# Antibacterial and Antiproliferative Properties of *Portulaca oleracea* Essential Oil and its Targeted Delivery to MCF-7 Cells via Spermine-PLA-PEG-FA Nanocapsules

**DOI:** 10.1101/2025.08.25.672136

**Authors:** Rasool Asghari Zakaria, Mehran Noruzpour, Shima Bourang, Hashem Yaghoubi

## Abstract

*Portulaca oleracea* essential oil is plant-derived product with documented antimicrobial and antiproliferative activities. Its clinical use is limited by poor water solubility, chemical instability, and lack of targeting. Encapsulation in polymeric nanocarriers can enhance solubility, stability, and selective delivery. This study investigated the antibacterial activity of *P. oleracea* essential oil against Staphylococcus aureus and Escherichia coli, and its antiproliferative effects on human breast cancer (MCF-7) cells using folic acid-modified spermine-polylactic acid-polyethylene glycol (Spermine-PLA-PEG-FA) nanocapsules. Gas chromatography–mass spectrometry (GC–MS) identified major components such as α-pinene, limonene, and phytol. The essential oil and quercetin were co-encapsulated in the nanocapsules. Characterization was conducted by proton nuclear magnetic resonance spectroscopy (¹H-NMR), Fourier transform infrared spectroscopy (FTIR), thermogravimetric analysis (TGA), differential thermogravimetric analysis (DTG), and transmission electron microscopy (TEM). Antibacterial properties were assessed using disk diffusion, minimum inhibitory concentration (MIC), and minimum bactericidal concentration (MBC) assays. Antioxidant capacity was measured via 2,2-diphenyl-1-picrylhydrazyl (DPPH) assay. Cytotoxicity was evaluated by 3-(4,5-dimethylthiazol-2-yl)-2,5-diphenyltetrazolium bromide (MTT) assay and flow cytometry. Nanocapsules had spherical shape, size of 115-237 nm, and pH-responsive release. Quercetin-loaded nanocapsules induced apoptosis in MCF-7 cells with an IC₅₀ of 11.21 mg/µl. This delivery system improved the antibacterial and anticancer efficacy of *P. oleracea* essential oil.

**Highlights:**

- The presence of medicinal, antibacterial and anti-proliferative compounds in *Portulaca oleracea* essential oil controls the growth of bacteria (both gram-positive and gram-negative strains), especially *Staphylococcus aureus bacteria*, which are representative of gram-positive bacteria.
- Encapsulating *P. oleracea* essential oil in Spermin-PLA-PEG-FA copolymer nanoparticles prevented its oxidation and destruction in free conditions without a polymer coating.
- The use of folic acid as a cell marker in the copolymer nanoparticle structure led to the identification of MCF-7 cancer cells by the nanoparticle/plant essential oil complex. This action increased the efficiency of the essential oil-targeted transfer system.
- The presence of compounds such as resveratrol and lycopene in *P. oleracea* essential oil has increased its antioxidant properties.

## Introduction

One of the deadliest diseases is cancer, which kills people every year. According to Arnold (Arnold et al., 2022) and Giaquinto et al.(Giaquinto et al., 2022), breast cancer ranks among the most prevalent cancers affecting women globally. The principal treatment options for this condition include surgery, chemotherapy, and radiotherapy (Bourang, Noruzpour, Jahanbakhsh Godekahriz, et al., 2024). Nonetheless, many chemotherapy agents frequently induce side effects that impact various healthy cells and tissues, thereby diminishing the overall effectiveness of the treatment (Oun et al., 2018; Zhang et al., 2018). As a result, cancer treatment may benefit from the use of low-risk and effective drugs that inhibit cancer cell proliferation without harming healthy individuals (Bourang, Asadian, et al., 2024). The therapeutic effects of many medicinal plant essential oils are still unknown because of their diverse medicinal properties (Asghari Zakaria et al., 2024).

*Portulaca oleracea* stands out as a significant medicinal plant, with numerous studies validating its efficacy in treating various conditions, including hyperglycemia and hyperlipidemia (Montoya-García et al., 2023; Srivastava et al., 2023). In addition, the extract and essential oil (EO) of *P. oleracea* have high potential to prevent bacterial and cancer cell proliferation (Ojah et al., 2021). Essential oils derived from medicinal plants have therapeutic properties; however, the use of EOs in their free forms may be limited by several critical factors, such as low stability and susceptibility to degradation through volatilization and/or oxidation when exposed to external deteriorating elements such as oxygen, light, and temperature; low water solubility; and potential interactions with food components (Sh et al., 2024). Hence, increasing the solubility of essential oils can increase their efficacy in treating cancer (Sh et al., 2024).

Nanocarriers are widely used to protect and deliver EOs more effectively than in their free form, largely owing to their ability to interact with cell receptors (Noruzpuor et al., 2024). Biomimetic nanotechnology, inspired by plant structures, enables the encapsulation of EOs via polymer-based systems such as nanoparticles, micelles, and nanocapsules (Chiriac et al., 2021). These carriers form through the entanglement of polymer chains, creating a protective matrix that stabilizes and traps EO droplets during delivery. EO carrier systems are often designed to mimic plant tissue structures and are influenced by factors such as plant species, extraction methods, EO chain length, and treatment conditions (Alshawwa et al., 2022). Polysaccharides, which are naturally abundant and biodegradable carbohydrate polymers, are commonly used because of their safety and effectiveness in encapsulating long-chain EOs and enabling controlled release (Cimino et al., 2021). Additionally, a plant’s developmental stage can significantly affect the choice of carrier system.

In addition to plant-inspired systems, various nanocarriers have been developed to improve the solubility and bioavailability of hydrophobic drugs (Lin & Chen, 2020). Micelles formed from amphiphilic copolymers, particularly polylactic acid–polyethylene glycol (PLA–PEG), are widely used because of their biodegradability, biocompatibility, and ability to self-assemble in aqueous environments (Bourang, Noruzpour, Azizi, et al., 2024). However, PLA-based nanoparticles can interact with plasma proteins, increasing the particle size and reducing the circulation time (Swetha et al., 2023). To overcome this, hydrophilic PEG chains are introduced to mask the surface and reduce recognition by the reticuloendothelial system (RES) (Yang et al., 2012). Spermine, a naturally occurring polyamine, has also been incorporated into these systems to increase cellular uptake and endosomal escape, especially for gene delivery (Amani et al., 2019). When conjugated to PLA–PEG copolymers, spermine introduces positive charges that facilitate electrostatic interactions with negatively charged nucleic acids or cell membranes (Subroto et al., 2023). This modification not only stabilizes the nanoparticle structure but also has potential for targeted and efficient delivery of both hydrophobic drugs and genetic material (Ahmadi-Nouraldinvand et al., 2024). According to Hong et al.(Hong et al., 2023), cancer cells exhibit a significant presence of folic acid receptors on their surfaces. These ligands can be used as a means for the targeted delivery of pharmaceutical agents to cancer cells (Martín-Sabroso et al., 2021; Salim et al., 2020). While this strategy has been well explored for conventional therapeutics, its application in the targeted delivery of EOs remains limited (Kita & Dittrich, 2011). The encapsulation of EOs into nano-or microscale carriers not only protects them from degradation, oxidation, and evaporation but also enhances their bioavailability and controlled release. These carriers allow EOs to diffuse more efficiently into target tissues, requiring lower doses while minimizing side effects, sensory impacts, and microbial resistance. Effective encapsulation and delivery depend on multiple factors, including carrier type, EO molecular size, surface ligands, and cellular conditions, all of which are especially relevant when working with complex, long-chain plant-derived compounds. (Bourang, Noruzpour, Jahanbakhsh Godekahriz, et al., 2024).

On the other hand, *P. oleracea* EO has demonstrated notable antibacterial activity against a variety of pathogenic bacteria (Tleubayeva et al., 2021). Studies have shown its effectiveness, particularly against both gram-positive bacteria such as *Staphylococcus aureus* and gram-negative bacteria such as *Escherichia coli* (Keser et al., 2021). These antimicrobial properties are attributed to the rich composition of bioactive compounds within the oil, including terpenoids, phenolics, and fatty acids, which can disrupt bacterial membranes and inhibit their growth. Key constituents such as α-pinene, limonene, and phytol are known to disrupt bacterial membranes and inhibit growth, highlighting the potential of oil as a natural antimicrobial agent, especially in the context of increasing antibiotic resistance (Fouda et al., 2022).

Despite the well-documented antibacterial and anticancer properties of *P. oleracea* EO, its practical use is limited by poor solubility, instability, and nonspecific distribution in biological environments. Although nanocarriers such as PLA-PEG copolymers have been widely studied for drug delivery, the integration of spermine for improved cellular uptake and folic acid for targeted delivery to folate receptor-expressing cancer cells remains underexplored, particularly for essential oils. Moreover, while the antimicrobial activity of *P. oleracea* EO against pathogens such as *Staphylococcus aureus* and *Escherichia coli* has been recognized, targeted nanodelivery systems that combine these antibacterial effects with anticancer applications have not been sufficiently investigated. This gap emphasizes the need for multifunctional nanocarriers that can simultaneously address antibacterial and anticancer challenges through enhanced targeted delivery of natural bioactive compounds.

This study aimed to evaluate the antibacterial activity of *Portulaca oleracea* EO against *Staphylococcus aureus* and *Escherichia coli*, as well as its antiproliferative effects on MCF-7 breast cancer cells. For anticancer evaluation, a targeted delivery system was developed using spermine-PLA-PEG nanocapsules functionalized with folic acid receptors on their surface, enhancing their selective uptake by cancer cells. This approach aims to improve the bioavailability, stability, and targeted delivery of EOs, minimizing systemic toxicity and maximizing therapeutic efficacy.

## Materials and methods

### Materials

In accordance with our previous study (Amani et al., 2021), the PLA-PEG-FA copolymer was synthesized. The following materials were purchased from Sigma‒Aldrich (USA): LB medium (Product No. 51208), RPMI 1640 (Product No. R2405), dimethyl sulfoxide (DMSO) (Product No. D8418), fetal bovine serum (FBS) (Product No. TMS-016), 3-(4,5-dimethyl-2-thiazolyl)-2,5-diphenyl-2H-tetrazolium bromide (MTT) (Product No. 475989), n-hexane (n-hexane basis) (Product No. 1.04394), and a penicillin/streptomycin solution (Product No. P7539). Strains of *Staphylococcus aureus* (ATCC 25923) and *Escherichia coli* (ATCC 25922) were acquired from Abosina (Iran), while the MCF-7 cell line (NCBI No. C135, human breast cancer cell line) was sourced from the Pasteur Institute of Iran. The Annexin V-FITC/propidium iodide (PI) Kit I was purchased from BD Biosciences Pharmingen (USA) (catalog no. BMS500FI-300), and acridine orange/ethidium bromide (AO/EB) (catalog no. L13159.06/17898) alongside 2,2-diphenyl-1-picrylhydrazyl (DPPH) (catalog no. 044150.03) was obtained from Thermo Fisher Scientific.

Preliminary research for this study began in 2021. Plant cultivation began in early 2022 and continued until late 2023, at which point all tests and measurements were completed.

### Plant material and explant source (*in vivo* production)

*P. oleracea* seeds with herbarium codes 001/001/151 were collected from arable lands in Ahvaz County (capital of Khuzestan Province), Iran. The purchased seeds were cultivated on the research farm of the University of Mohaghegh Ardabili. After the *P. oleracea* aerial parts were collected, healthy, fresh and phenotypically suitable plants were washed with running water and then dried for 72 hours at room temperature (25°C) in the dark. After the leaf samples were dried, they were cut into smaller pieces and prepared for essential oil extraction (Montoya-García et al., 2023).

### Methods for extracting essential oils from the aerial parts of *P. oleracea*

The powder of each sample (30 g) was accurately measured, placed in a sealed container with 150 ml of water, and then left to sit for 30 minutes. Afterwards, the mixture was transferred to a Clevenger apparatus flask, and 1.5 ml of n-hexane was added to the extraction machine’s measuring tube before boiling for 5 hours. The extract was dried over anhydrous sodium sulfate and then diluted to 2.0 ml with n-hexane (Zhu et al., 2010).

### Identification and quantification of the essential oil components

The essential oil of *P. oleracea* was analyzed via GC-MS (Agilent 7890B series GC‒MS). The components were identified by matching their mass spectra to the Wiley Registry 9th Edition/NIST 2011 mass spectral library and comparing their retention indices to known compounds or values from the literature (Rahim et al., 2023). The relative percentages of these components were determined by electronically integrating the peak areas without using a correction factor.

### Disk Diffusion Assay

This research investigated the antimicrobial effects of *P. oleracea* essential oil using *Staphylococcus aureus* (ATCC 25923) as a model for gram-positive bacteria and *Escherichia coli* (ATCC 25922), which are gram-negative bacteria, in three replicates. A colony of each bacterial strain was introduced into 50 mL of LB medium (25 g/L) and incubated at 37°C with continuous shaking at 45 rpm for 12 hours. Following this incubation period, the cultures were diluted to a 0.5 McFarland standard and spread across a Petri dish containing 20 mL of LB agar medium (40 g/L). Various volumes of *P. oleracea* essential oil (1, 2, 5, 10, 15, and 20 µL) were applied to sterile disks measuring 6 mm in thickness, which were subsequently placed on agar plates after they were allowed to dry (Donaldson et al., 2005). The plates were then incubated at 37°C for 48 hours, after which the diameters of the inhibition zones were measured via ImageJ software.

### MIC and MBC determination

By using the serial microplate dilution method, various concentrations of *P. oleracea* essential oil (200, 100, 50, 25, 12.5, 6.25, 3.125, and 1.56 µL) were added to each well of the 96-well plates for MIC determination (six replicates for reaching the bacterial strain). After standardizing the *E. coli* and *S. aureus* suspensions to a minimum McFarland concentration, 20 mL of each bacterium was added to each well of a 96-well plate and incubated at 37°C. The bacteria were observed after 48 h, and the concentration of *P. oleracea* essential oil that inhibited the growth of the bacteria was designated the MIC. To determine the MBC, 50 µL of bacteria from the nongrowing wells (the wells treated with *P. oleracea* essential oil with clear media) were spared on the surface of the LB agar medium via soap and incubated for 48 h at 37°C (Janani et al., 2019).

### DPPH Assay

The antioxidant properties of *P. oleracea* essential oil were determined on the basis of its ability to scavenge DPPH radicals in three replicates. The essential oil of *P. oleracea* contains biochemical compounds with antioxidant properties. These biochemical compounds are considered the basis of the present research hypothesis. Consequently, a modified version of the methodology outlined in prior studies was employed to assess the antioxidant properties of *P. oleracea* essential oil. Specifically, 1 mL of a 0.5 mM DPPH solution in methanol was mixed with 3 mL of a methanol solution containing varying concentrations of *P. oleracea* essential oil (10, 25, 50, and 100 μL mL^-1^). The samples were incubated in the dark at 25°C for 30 minutes, after which their absorbance was measured at 520 nm (Tsimogiannis et al., 2017). The percentage of DPPH radical scavenging was ultimately calculated via the following equation (Formula 1):

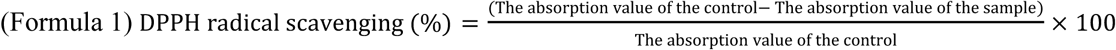

Note that as a control, 3 mL of methanol containing 0.5 mM DPPH was used in this experiment.

### Synthesis of PLA-PEG-FA nanoparticles

For this purpose, 0.2 g of folic acid (441.4 molecular weight Da) was first dissolved in 10 ml of DMSO and activated by adding 0.1 g of EDC and 0.06 g of NHS for 2 h at room temperature under nitrogen. This step prepares the carboxyl group of folic acid for reaction with the terminal amine of PEG. Then, 0.5 g of polyethylene glycol amino acid (PEG-NH₂ with a molecular weight of 5,000 Da) was added to the activated FA solution and stirred for 24 h in the dark at room temperature. The PEG-FA product was purified by precipitation in cold ether and centrifugation (Eppendorf, Germany) at 10,000 rpm for 15 min, and after washing with ethanol, it was dried via a freeze-dryer (Thermo Fisher Scientific Co., USA). Then, 1 g of polylactic acid (PLA-COOH with a molecular weight of 10.000 Da) and 0.6 g of PEG-FA were reacted in the presence of 0.15 g of DCC and 0.075 g of DMAP in 30 ml of DMSO for 24 h. The PEG-FA-PLA copolymer was purified by dialysis against distilled water for 48 h and collected by lyophilization (**Figure 1**).

**Figure 1.**
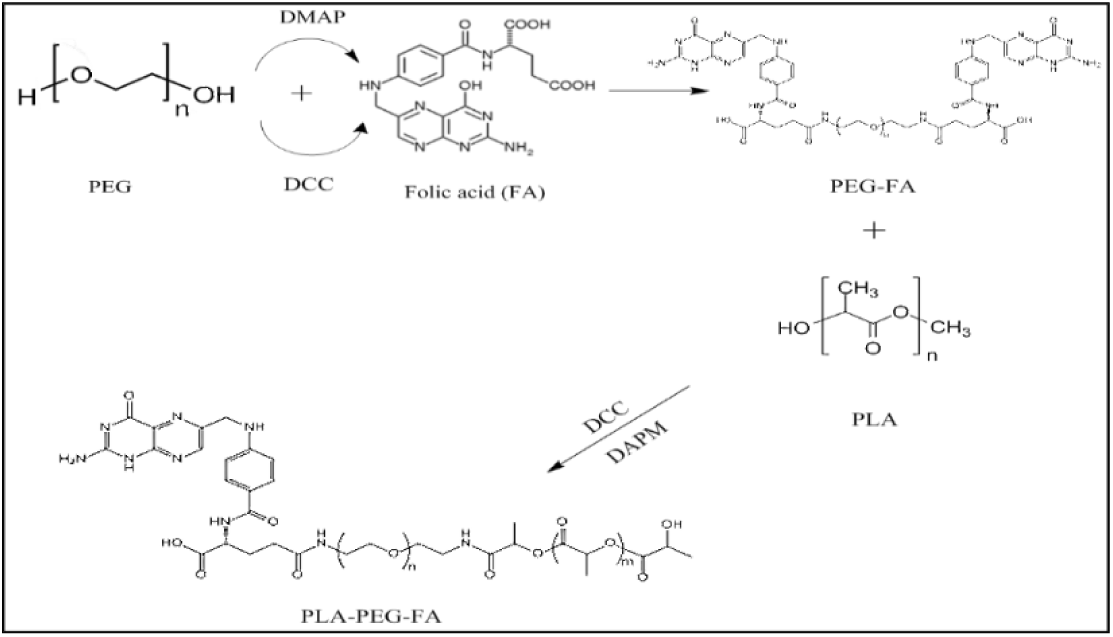
PLA-PEG-FA nanoparticle synthesis steps.

### Synthesis of Spermine-PLA-PEG-FA (SPPF) nanoparticles

For this purpose, first, 0.4 g of PEG-FA-PLA nanoparticles were reacted with 0.08 g of spermine in the presence of 0.05 g of EDC and 0.03 g of NHS in 20 ml of DMSO for 24 h, and the final product, SPPF, was purified by dialysis against PBS (pH=7.4) for 72 h (**Figure 2**). Then, 10 mg of PEG-FA-PLA-spermine was dissolved in 1 ml of acetone and added to 10 ml of water containing 0.1% Polyvinyl alcohol (PVA) with stirring at 1,000 rpm. After evaporation of the acetone, the nanoparticles were collected via centrifugation (15,000 rpm, 20 min) and washed with distilled water.

**Figure 2.**
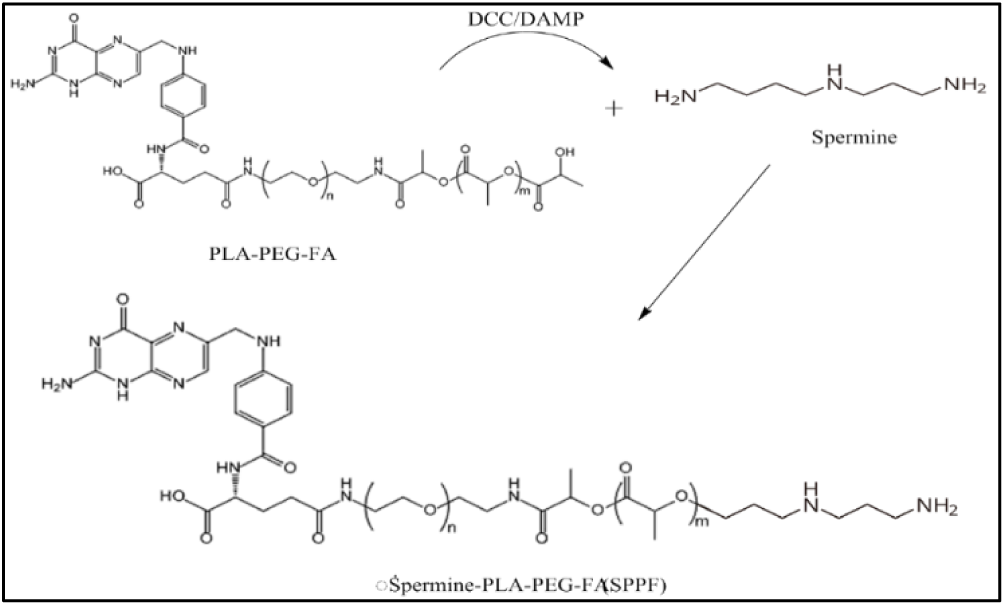
Spermine-PLA-PEG-FA (SPPF) nanoparticle synthesis steps.

### Characterization of the nanoparticles

Infrared spectroscopy (FTIR-ABB Bomem-MB) and hydrogen nuclear magnetic resonance spectroscopy (^1^H-NMR-Bruker 400 MHz) were used to analyze the produced nanoparticles. Dynamic light scattering (DLS, Malvern Instruments, Westborough, MA, USA) was used to analyze the nanoparticle shape.

### Synthesis of Spermine-PLA-PEG-FA/Essential Oil (SPPFE) or Quercetin (SPPFQ) Nanocapsules

For this purpose, the method of Mohajeri et al. 2025 was used with some modifications. First, 500 μL of Portulaca oleracea essential oil or 500 μL of quercetin solution (control sample) was added separately to 1 mL of chloroform containing 50 mg of SPPF nanoparticles and sonicated on ice for 30 seconds. Then, the resulting mixture was emulsified in 3 mL of water containing 0.3% (w/v) PVA under sonication for 2 min. The chloroform in the mixture was completely evaporated via a rotary evaporator (Heidolph Co., Germany), and the resulting nanoparticles were collected, washed with 100 mL of phosphate buffer and centrifuged (12,000 rpm for 20 min). Then, 10 mL of phosphate buffer was added to the resulting nanoparticles and filtered through a 0.45 μm membrane filter to remove large nanoparticles. Finally, the nanoparticles were dried via a freeze-dryer and stored at –80°C (Mohajeri, Yaghoubi, et al., 2025).

### Characterization of Nanocapsules

Transmission electron microscopy (TEM, JEM-2100) and dynamic light scattering (DLS, Malvern Instruments, Westborough, MA, USA) were used to analyze the nanocapsule shape, particle size, and zeta potential, respectively.

### Kinetics of the Release of Essential Oil (EO) or Quercetin (Que) from Nanocapsules

The release of EO or Que from SPPFE or SPPFQ nanocapsules was evaluated. Owing to the different pH levels in normal and cancer tissues, release assays were performed in two separate environments under acidic (pH=5.5) and neutral (pH=7.4) conditions. The nanocapsules were incubated in 20 mL of PBS at 37°C. At scheduled time intervals (ranging from 15 min to 72 h), the nanocapsules were collected by centrifugation (16602 × g for 30 min), and the resulting supernatant was used for release assays. The nanocapsules were then resuspended in fresh buffer and incubated for the next time interval (Bourang, Noruzpour, Azizi, et al., 2024). A NanoDrop spectrophotometer with a wavelength of 370 nm was used to measure the concentration of each compound in each sample. The cumulative percentage of EO or Que released from the nanocapsules was calculated via the following formula:

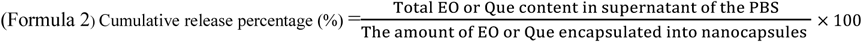

### Cell culture

We obtained the MCF-7 human breast cancer cell line (ATCC^®^-HTB-22) from the Pasteur Institute’s National Cell Bank of Iran. In a humidified incubator set at 37°C with 5% CO_2_, the cells were cultivated in RPMI 1640 (Gibco, USA) supplemented with 10% heat-inactivated FBS (Gibco, USA). The cells were passaged once they reached 70–80% confluence via trypsin-EDTA (Gibco, USA) to support continuous exponential growth (Jahazi & Akbari, 2020).

### *In vitro* cytotoxicity studies

The MTT assay is a commonly used technique for assessing cytotoxicity. It operates on the principle that viable cells convert MTT into insoluble formazan crystals, reflecting their mitochondrial activity. Since mitochondrial function typically halts upon cell death, the MTT assay serves as a reliable indicator of cell viability (Constante et al., 2022). In this study, the cytotoxic effects of purslane essential oil, quercetin (at concentrations of 0, 5, 10, 25, 50, and 100 μL/mL) and SPPF, SPPPFE, and SPPFQ nanocapsules (at concentrations of 0, 5, 10, 25, 50, and 100 μg/mL) were evaluated via the MTT assay. MCF-7 cells were seeded in 96-well plates with 200 μL of complete medium (RPMI 1640 with 10% FBS) at a density of 7 × 10^3^ cells per well. The cells were incubated for a full day in an environment with 5% CO_2_ at 37°C. For an additional twenty-four hours at the same temperature, the samples were then exposed to varying concentrations of nanocapsules (0 to 100 µg/mL). The MTT assay and a BioTek Instruments reader (Winooski, VT, USA) were used to measure the absorbance at 570 nm, and the cell viability was evaluated. The percentage of surviving cells was then calculated via the following equation (Bourang, Asadian, et al., 2024):

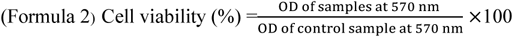

### Cell Apoptosis Analysis

One of the biomarkers of the apoptosis process is the translocation of phosphatidylserine from the inner plasma membrane to the outer plasma membrane. When phosphatidylserine is present on the cell surface, it can be detected by fluorescence staining (Amani et al., 2021), which results in the binding of phosphatidylserine to annexin. Notably, annexin is used to detect phosphatidylserine present on the outer cell membrane, whereas the combination of annexin and propidium iodide is used to distinguish between apoptotic and necrotic cells.

According to the manufacturer’s instructions, flow cytometry analysis was performed via an Annexin V-Dy634 kit (Annexin V Apoptosis Detection Kit with PI, Immunostep) to investigate the apoptotic response of MCF-7 cells to concentrations obtained from the IC_50_ test of quercetin and *P. oleracea* essential oil (in free form) and SPPFE and SPPFQ nanocapsules. Untreated cells were used as positive controls. FlowJoTM software (version 10; FlowJo LLC) and a flow cytometer (FACS Verse, BD Bioscience) were used for data collection and analysis (Bourang et al., 2025).

### Statistical analysis

Each quantitative parameter investigated in this study was measured in at least three replicates. The data were analyzed via one-way ANOVA with SPSS software. For mean comparisons, Duncan’s multiple-range test was applied at a five percent significance level, and the Kolmogorov‒Smirnov test was used to determine the normality of the data set. The results of each treatment are presented as the mean ± standard deviation (mean ± SD).

## Results

### GC‒MS analysis of *P. oleracea* essential oils

According to the results from the analysis of the mass of the essential oil of *P. oleracea* (**Figure 3** and **Table 1**), the presence of compounds such as 2-Pentyl-furan, isoeugenol, humulene, cedrol, myristic acid, and pentatriacontane was associated with anti-inflammatory, antibacterial, antioxidant, antidiabetic, and anticancer properties.

**Figure 3.**
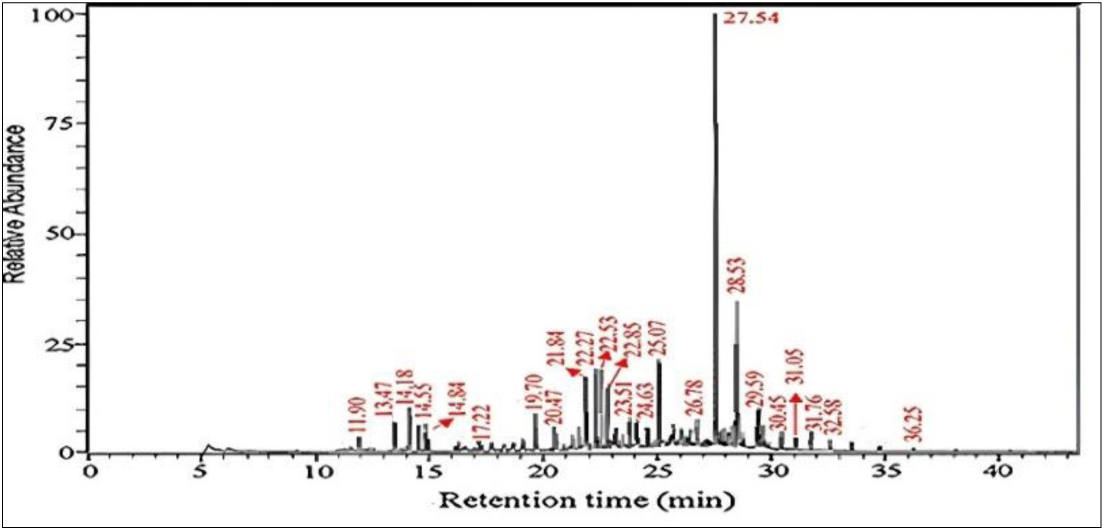
Schematic image of the results obtained from *P. oleracea* essential oil GC‒mass spectrometry.

**Table 1.** Results of the GC-mass analysis of the essential oil of *P. oleracea*.

| Compound | Formula | Rt (min) | Biological properties | Ref |
| --- | --- | --- | --- | --- |
| 2-Pentyl-furan | C<sub>9</sub> H<sub>14</sub>O | 11.90 | Antioxidant, Antitumor, Antibacterial and Antifungal | )Bhandari et al., 2011; Tian et al., 2022; Zhang et al., 2024( |
| Benzene acetaldehyde | C<sub>8</sub> H<sub>8</sub>O | 13.47 | Antitumor and Antibacterial | (Sri et al., 2021; Tadigiri et al., 2020) |
| Cis-Linalool oxide | C<sub>10</sub> H<sub>18</sub>O<sub>2</sub> | 14.18 | Antioxidant and Antibacterial | (Zhao et al., 2024) |
| Cyclopropane Ethanol | C<sub>8</sub>H<sub>14</sub>O | 14.55 | Antioxidant, Antitumor, Antibacterial and Antifungal | )Barreiro-Costa et al., 2021; Duanis-Assaf et al., 2023; Karlinsey et al., 2022( |
| Santolina triene | C<sub>10</sub>H<sub>16</sub> | 14.84 | Antitumor and Antibacterial | (Khammassi et al., 2024; Saygideger et al., 2021) |
| 3-Methyl-6-(1-methylethylidene)-cyclohexene | C<sub>10</sub>H<sub>16</sub>O | 17.22 | Antioxidant, Antitumor and Antibacterial | (Selvaraju et al., 2021) |
| 2-Methoxy-4-vinylphenol | C<sub>9</sub>H<sub>10</sub>O<sub>2</sub> | 19.70 | Antioxidant and Anti-inflammatory | (Asami et al., 2023; Rubab et al., 2020) |
| Isoeugenol | C<sub>10</sub>H<sub>12</sub>O<sub>2</sub> | 20.47 | Antioxidant, Antitumor, Antibacterial and Antifungal | (Fernandes Melo Reis et al., 2024; Freitas, 2023; Goswami et al., 2022) |
| Alloaromadendrene | C<sub>15</sub>H<sub>24</sub> | 21.84 | Anti-Diabetic and Anti-inflammatory | (Barros et al., 2022; Tuan et al., 2021) |
| 4,11,11-Trimethyl-8-methylene-bicyclo-undec-4-ene | C<sub>15</sub>H<sub>24</sub> | 22.27 | - | - |
| Humulene | C<sub>15</sub>H<sub>24</sub>O | 22.53 | Anti-neuroinflammatory and Anti-inflammatory | (Cordeiro et al., 2023; Jang et al., 2020) |
| (1,2,6,7)-Tricyclo[5.3.1.1]dodecane-11,12-dione | C<sub>12</sub>H<sub>16</sub>O<sub>2</sub> | 22.85 | - | - |
| Decahydro-1,6-bis(methylene)-4-(1-methylethyl)-naphthalene | C<sub>15</sub>H<sub>24</sub> | 23.51 | - | - |
| Cis-Lanceol | C<sub>15</sub>H<sub>24</sub>O | 24.63 | Antimicrobial | (Thuy et al., 2021) |
| Cedrol | C<sub>15</sub>H<sub>26</sub>O | 25.07 | Anticancer, Anti-inflammatory, Anti-arthritic and Antioxidant | (Yun et al., 2022; Zhao et al., 2023) |
| Myristic acid | C<sub>14</sub>H<sub>28</sub>O<sub>2</sub> | 26.78 | Anti-fungal, Anti-viral, Anti-inflammatory, Antinociceptive, Anticancer, Anti-parasitic, and Immunomodulating activities | (Alonso-Castro et al., 2022; Javid et al., 2020) |
| 9-Eicosyne | C<sub>20</sub>H<sub>38</sub> | 27.54 | - | - |
| n-Hexadecanoic acid | C<sub>16</sub>H<sub>32</sub>O<sub>2</sub> | 28.53 | Antioxidant, Antitumor, Antibacterial and Antifungal | (Purushothaman et al., 2024) |
| 1-Mono-linolenin | C<sub>21</sub>H<sub>36</sub>O<sub>4</sub> | 29.59 | - | - |
| Heptacosane | C<sub>27</sub>H<sub>56</sub> | 30.45 | Antibacterial and Antifungal activities | (Al Kamaly et al., 2024) |
| Docosane | C<sub>22</sub>H<sub>46</sub> | 31.05 | Anti-inflammatory | (CUI et al., 2022) |
| Pentatriacontane | C<sub>35</sub>H<sub>72</sub> | 31.76 | Antimicrobial, Anti-biofilm, Antioxidant, Anti-parasitic, Antinociceptive, Anti-inflammatory, and Anticancer | (Roopa et al., 2020) |
| Dotriacontane | C<sub>32</sub>H<sub>66</sub> | 32.58 | Anti-inflammatory, Anti-androgenic, Anticancer, Antioxidant, Anti-inflammatory | (Manhas & Kaul, 2024) |
| Nonacosane | C<sub>29</sub>H<sub>60</sub> | 36.25 | Anti-inflammatory and Antioxidant | (Al Kamaly et al., 2024) |

## Antibacterial Assays

### Disk Diffusion Assay

The antimicrobial properties of *P. oleracea* essential oil were evaluated via disk diffusion, MIC, and MBC assays. The concentration of *P. oleracea* essential oil influenced the diameter of the growth inhibition zone. As shown in **Figure 4 A**, the growth inhibition zone diameter increased with increasing *P. oleracea* essential oil concentration. When the essential oil concentration was increased from 2 to 20 μL, the growth inhibition zone of *P. oleracea* against *S. aureus* increased in diameter from 7.6 to 14.2 mm. A similar trend was also observed for *E. coli* (**Figure 4 B**). The results revealed that *P. oleracea* essential oil was more effective against *S. aureus* than against *E. coli*.

**Figure 4.**
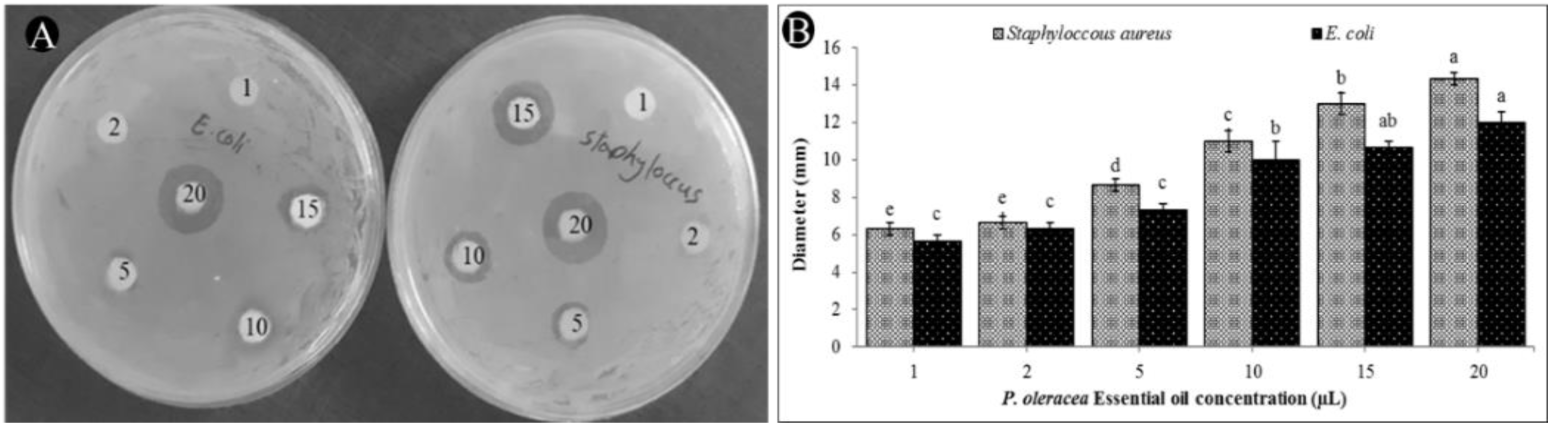
Antibacterial activity of *P. oleracea* essential oil; inhibition of *S. aureus* and *E. coli* by different concentrations of *P. oleracea* essential oil. (**A**), growth inhibition zone diameter of *P. oleracea* essential oil against *S. aureus* and *E. coli* (**B**). Nonsimilar letters indicate a statistically significant (*p* < 0.01) difference between different *P. oleracea* essential oil concentrations in each bacterial type.

### MIC and MBC determination

The MIC and MBC results revealed that *S. aureus* was less resistant to *P. oleracea* essential oil than was *E. coli*. The MIC and MBC values for *S. aureus* were 12.5 and 6.25 μL.mL^-1^, respectively, while these values were 25 and 12.5 μL.mL^-1^ for *E. coli* (**Figure 5**).

**Figure 5.**
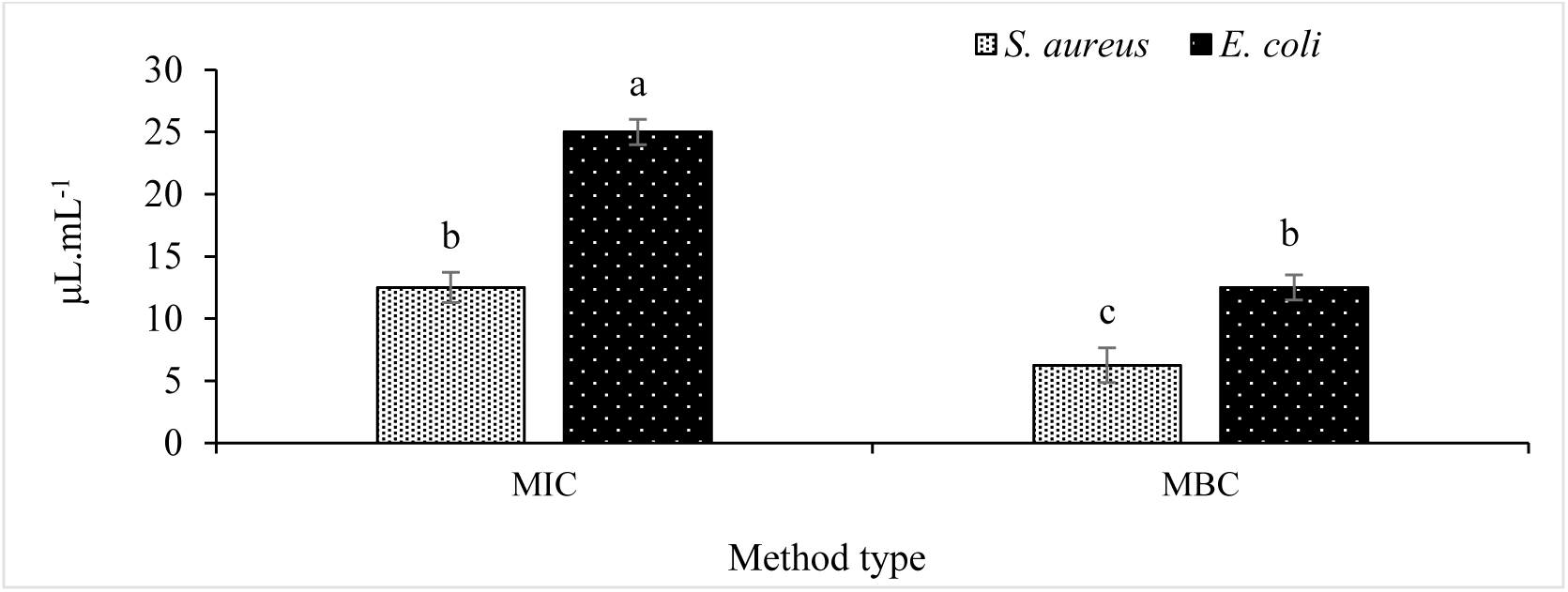
Antibacterial activity of *P. oleracea* essential oil; MIC and MBC concentrations of *P. oleracea* essential oil in controlling *S. aureus* and *E. coli* bacteria Nonsimilar letters indicate a statistically significant (*p* < 0.01) difference between different *P. oleracea* essential oil concentrations in each bacterial type.

### DPPH Assay

In this study, the antioxidant properties of *P. oleracea* essential oils were determined via the DPPH assay. The findings revealed that as the concentration of *P. oleracea* essential oil increased, so did the antioxidant activity (**Figure 6**). At a concentration of 100 μL.mL^-1^, the percentage of DPPH radical scavenging was the highest (27.61%). The concentration corresponding to 50% inhibition of the radical DPPH was 170.5 μL mL^-1^. As mentioned above, the reason for the antioxidant activity of *P. oleracea* essential oil is the presence of biochemical compounds with antioxidant properties, which have been identified via the GC‒mass spectrometry technique. The results of this technique are given in **Table 1** (Supplementary Data).

**Figure 6.**
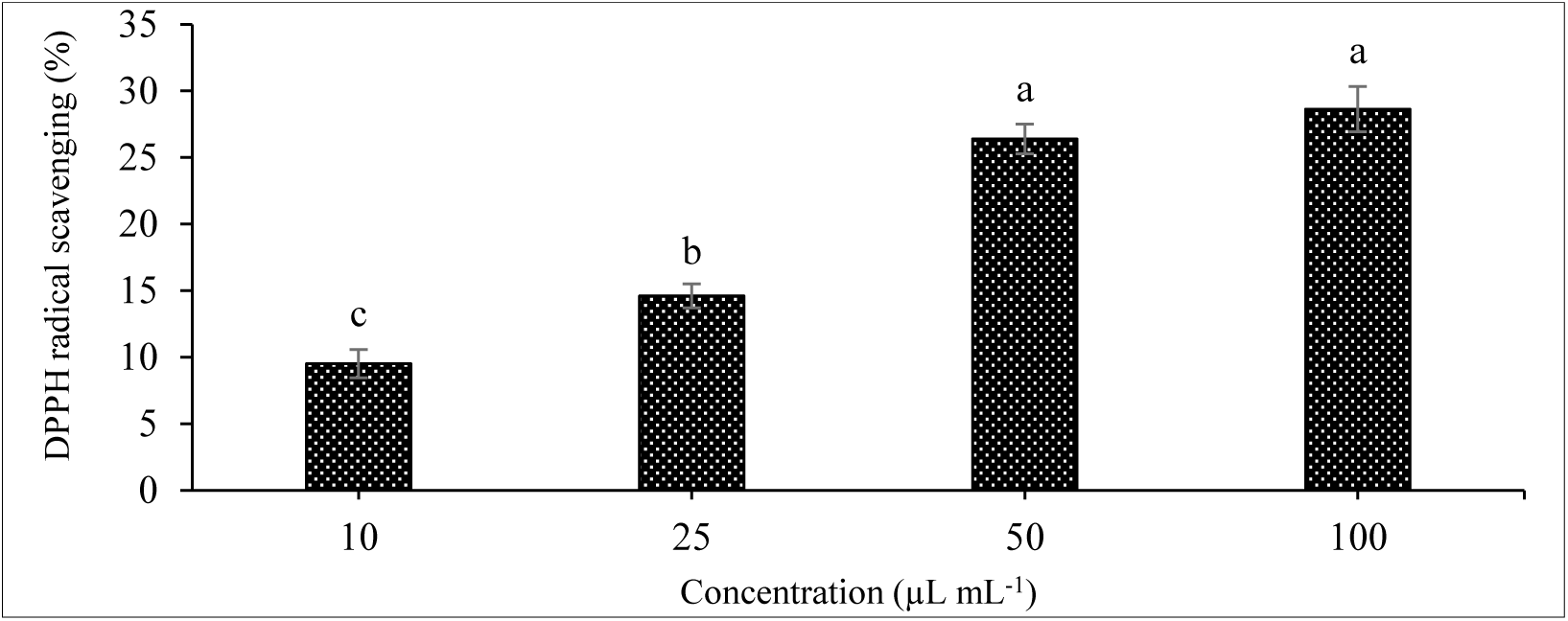
Antioxidant activity of different concentrations of *P. oleracea* essential oils determined via the DPPH assay. Nonsimilar letters indicate a statistically significant (*p* < 0.01) difference between different concentrations.

### Investigation of the structure of SPPF nanoparticles

#### 1H-NMR

The results of the ^1^H-NMR measurement of the SPPF copolymer revealed that the peaks at 1.7 and 2.4 ppm were related to the N-H and NH_2_ hydrogens present in spermine (**Figure 7 No-1**). In addition, the peak observed at 2.1 ppm was related to the carbon chain and N-H hydrogens present in PLA (**Figure 7 No-2**). Additionally, peaks related to the hydrogen atom in the N‒H bond and OH in the main chain of PEG were observed at 2.8 and 3.7 ppm, respectively (**Figure 7 No-3** and **4**). In addition, peaks related to folic acid were observed in the range of 6 to 8 ppm (**Figure 7**).

**Figure 7.**
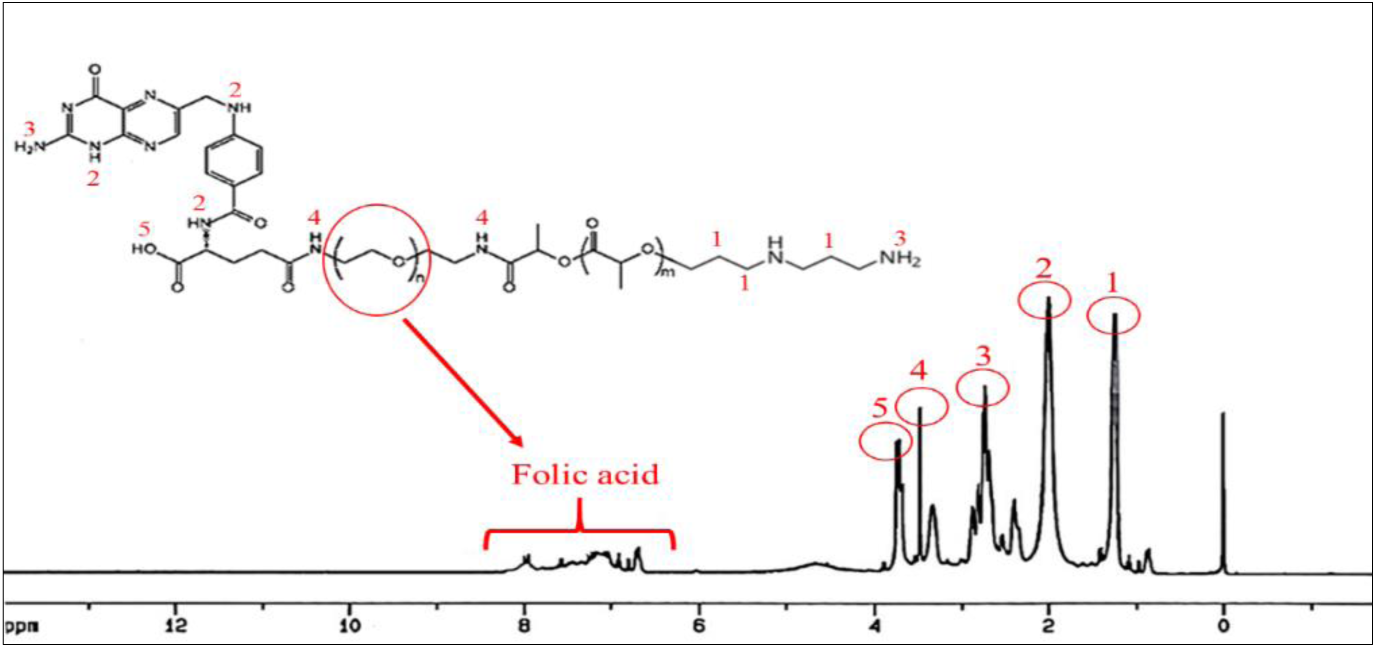
1H-NMR spectroscopy of spermine-PLA-PEG-FA (SPPF).

#### FTIR

The FTIR spectrum of PLA shows specific absorption bands that indicate its molecular structure, and the peak at 1750-1740 cm^-1^ corresponds to the carbonyl ester (C=O), which has a polymer effect. The additional peaks at 2950-2850 cm^-1^ are related to the C–H vibrations of the methyl and methylene groups, while the bands at 1450-1380 cm^-1^ reflect the C–H bending modes of the CH_3_ groups, and the regions at 1300-1080 cm^-1^ indicate the C– O ester (**Figure 8 A**). The FTIR spectrum of PEG (**Figure 8 B**) shows distinct absorption bands characteristic of its polyether structure, the most prominent of which is the strong O-H stretching vibration at 2870 cm^-1^. There is a sharp and intense peak at 1150 cm^-1^ dominating the spectrum, which is related to the stretching of the C-O-C ether bond that forms the PEG backbone. The bending vibrations of the CH_2_ groups are observed as bands of moderate intensity at 1540 cm^-1^. There are also sharp peaks in the range of 1350-1250 cm^-1^, which are related to torsional motions and C-H vibrations. The FTIR results of spermine (**Figure 8 C**) revealed the presence of amine groups and carbon chains. The broad peak at 3470 cm^-1^ is related to N‒H stretching in primary (NH_2_) and secondary (NH) amine groups. A sharp peak due to C-H stretching in methylene (CH_2_) groups was observed in the range of 2850 cm^-1^. The region of 1600–1650 cm^-1^ is related to N–H deformation in primary amines, whereas the sharp peak at 1530 cm^-1^ indicates N–H deformation in secondary amines. The sharp peaks in the range of 1150 cm^-1^ are related to C–N stretching. Additionally, the sharp peak in the range of 1470 cm^-1^ indicates CH2 bending deformation (**Figure 8 C**). The FTIR spectrum of the spermine-PLA-PEG-FA (SPPF) nanoparticles revealed the structural features of all four components. The broad peak in the range of 3300-3500 cm^-1^ is related to the N–H stretching vibrations of the spermine groups. Methylene C–H stretches are observed at approximately 2900 cm^-1^ from all four components of the nanoparticles. The presence of a clear peak at 1750 cm^-1^ indicates the carbonyl C=O group in the folic acid structure, which appears as a sharp peak. In the region of 1650 cm^-1^, the aromatic C=C bending peak of folic acid is located. The intense peak in the range of 1100-1200 cm^-1^ is related to the C– O–C stretching vibrations in the PEG structure, which is one of its prominent features. The peaks obtained in the range of 1500-1450 cm^-1^ are related to the N–H structural change as well as the nitrogen-containing rings in folic acid (**Figure 8 D**).

**Figure 8.**
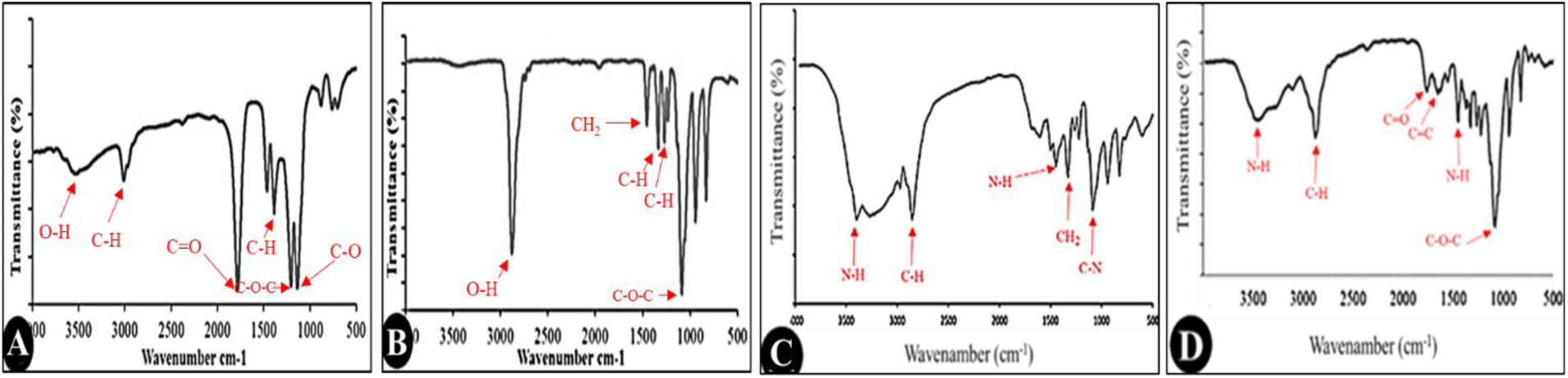
FT-IR spectra of **A)** PLA, **B)** PEG, **C)** spermine, and **D)** spermine-PLA-PEG-FA (SPPF)

#### TGA and DTG

The TGA and DTG results are shown in **Figure 9**, which presents the thermal degradation profiles of spermine, PLA-PEG-FA, and SPPF. The TGA curves (**Figure A, B, C**) show distinct degradation patterns. The results revealed that spermine decomposes rapidly at temperatures of approximately 200–300°C, indicating low temperature stability (**Figure 9 A**), whereas PLA-PEG-FA decomposes in two stages, indicating the presence of the three components FA, PEG, and PLA (**Figure 9 B**). The TGA results of the SPPF nanoparticles also indicate the high stability of the nanoparticles and their successful synthesis (**Figure 9 C**).

**Figure 9.**
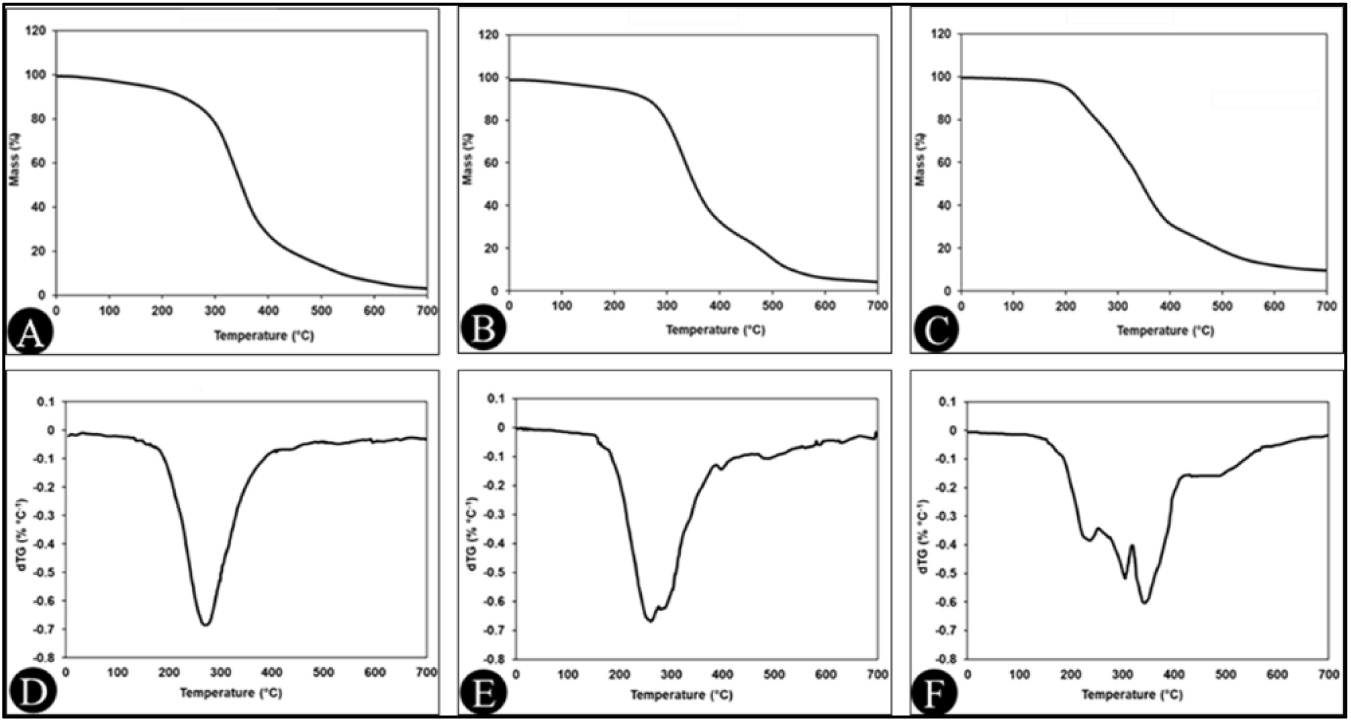
TGA (**A**, **B**, and **C**) and DTG (**D**, **E**, and **F**) curves for spermine, PLA-PEG-FA, and spermine-PLA-PEG-FA (SPPF), respectively.

The DTG curves characterize the degradation kinetics of the compounds over time and allow for the distinction between multicomponent and single-component compounds on the basis of the peaks presented therein. In **Figure 9 D**, the presence of only one sharp peak at 250°C indicates the presence of only one compound (spermine). This occurred while the PLA-PEG-FA nanoparticles were degraded in several steps (**Figure 9 E**). In addition, another multiple-step degradation of SPPF nanoparticles confirmed its successful synthesis (**Figure 9 F**).

### Investigation of the structure of SPPFE and SPPFQ nanocapsules

#### TEM

TEM images (**Figure 10**) showing the morphological characteristics of the SPPFE (**A**) and SPPFQ (**B**) nanocapsules. Both images show spherical-shaped nanoparticles, which is typical for polymer nanoparticles prepared via emulsion-based methods. The scale of 200 nm indicates that the nanoparticles are in the nanoscale range and have a relatively uniform size distribution. The presence of a dark core in the nanoparticles indicates the successful encapsulation of the essential oil in the polymer matrix. Compared with the SPPFE nanocapsules, the SPPFQ nanocapsules appeared to have a slightly more defined and consistent morphology, which could be related to the addition of quercetin instead of the plant essential oil. The presence of plant compounds with multiple active groups provides nanoparticles with a change in shape and polyhedrality. The absence of significant aggregation in both samples indicates effective stabilization by the PEG moieties. The uniform contrast in the images indicates the homogeneous distribution of essential oil in the nanoparticles. Overall, the TEM results confirmed the successful formation of nanoscale carriers with suitable dispersion and essential oil transport potential (**Figure 10 A** and **B**).

**Figure 10.**
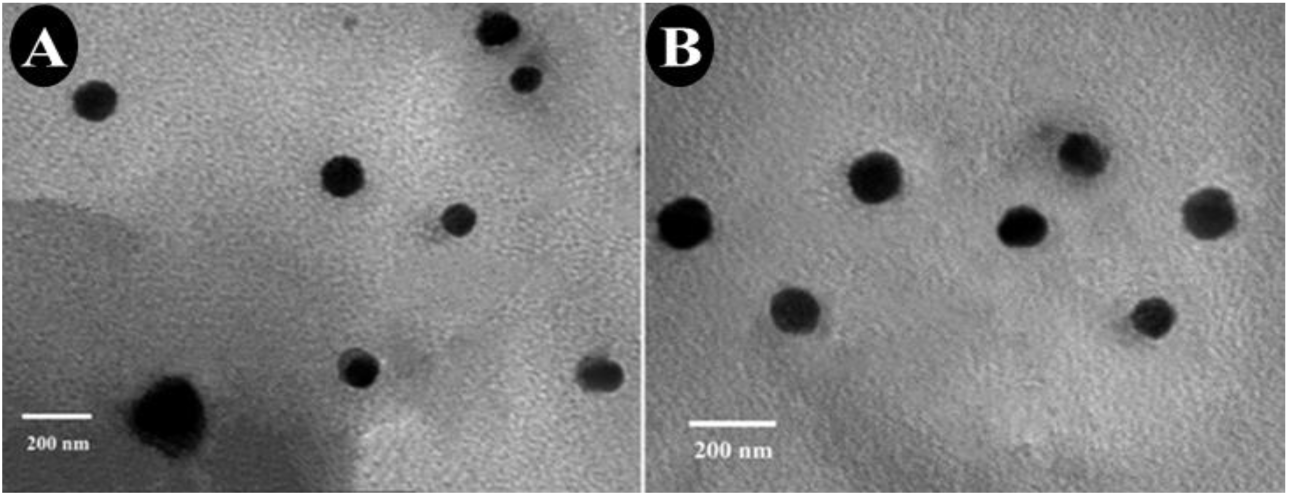
TEM images of SPPFE (**A**) and SPPFQ (**B**) nanocapsules.

#### DLS

The data in **Table 2** and **Figure 11** show a comprehensive analysis of the size distributions, zeta potentials, and dispersion indices (PDIs) of the SPPFQ and SPPFE nanocapsules. The size distribution plots (**Figure 11 A** and **C**) show that both types of nanoparticles are in the nanoscale range, with SPPFQ averaging 115 (±12) nm in size and SPPFE averaging 137 (±17) nm in size. The larger size of SPPFE than of SPPFQ could be related to the large size of the plant compounds present in the purslane essential oil, which contributes to the increase in the average particle diameter.

**Figure 11.**
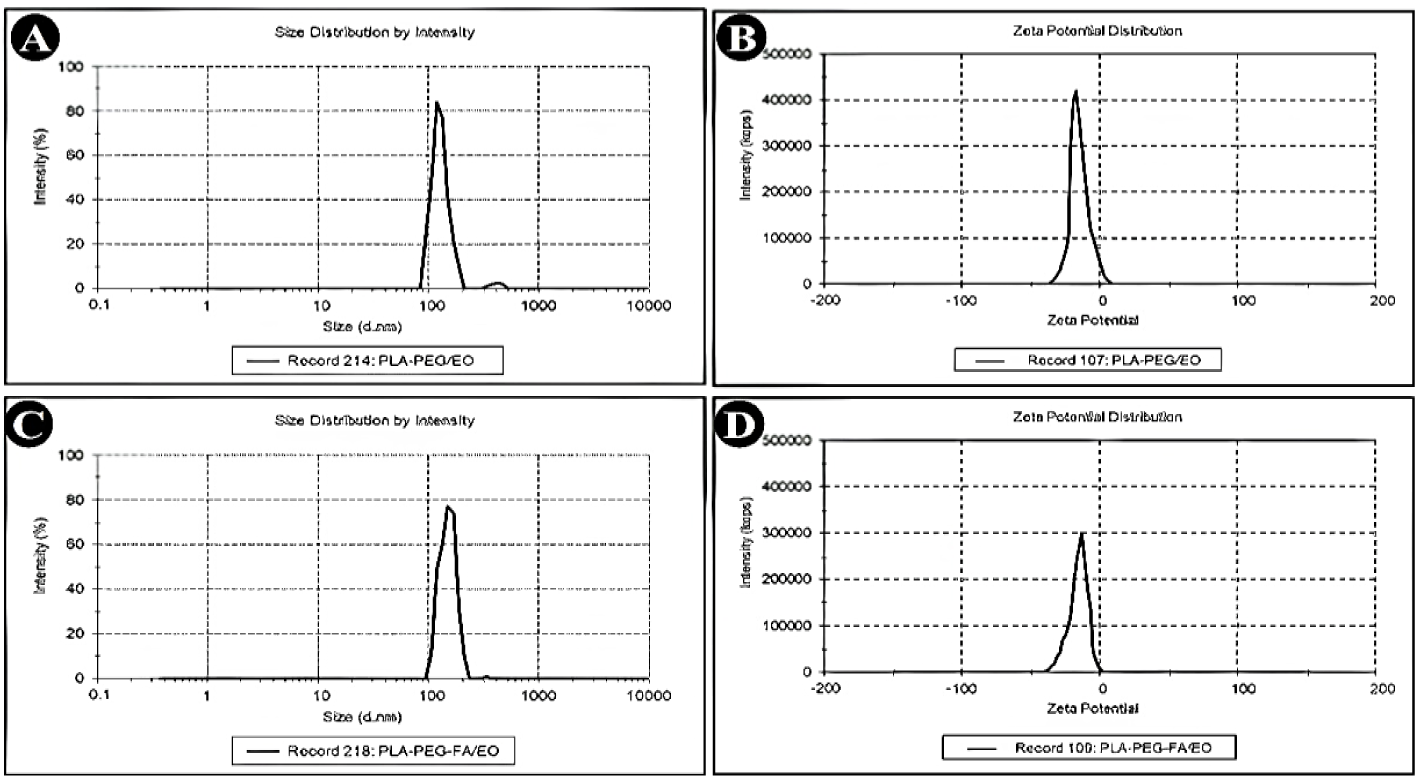
Particle size and zeta potential of the SPPFQ (**A** and **B**) and SPPFE nanocapsules (**C** and **D**).

**Table 2.**
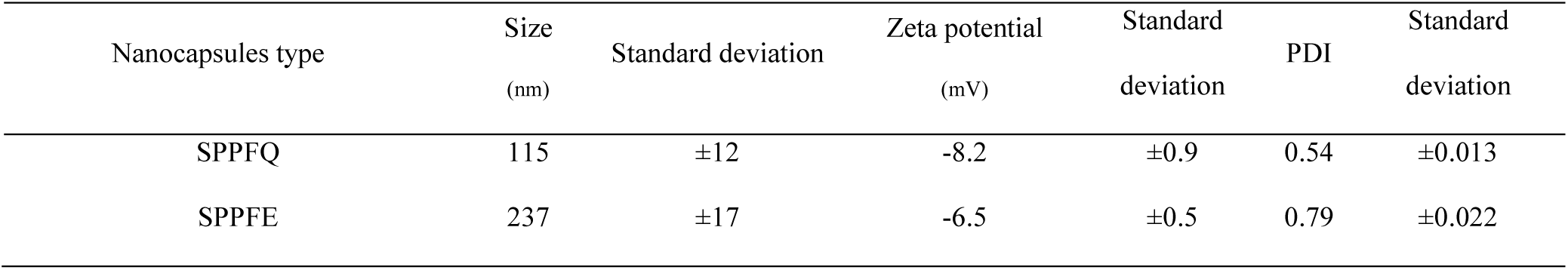
Size and zeta potential of the SPPFQ and SPPFE nanocapsules.

The Polydispersity Index (PDI) is a critical parameter describing the molecular mass distribution and size uniformity of nanoparticles. Lower PDI values indicate a more consistent particle size distribution, whereas higher values suggest polydisperse samples (Bourang et al., 2025). The PDI results for the nanoparticles examined in this work are provided as follows:

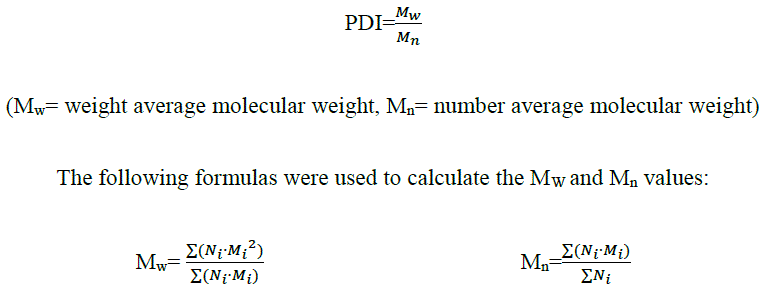

where N_i_ is the number of particles and M_i_ represents the mass of the particles.

The mean diameter and PDI are essential parameters as they determine the transport of nanoparticles *in vivo*. The PDI value can vary from 0.01 (monodisperse particles) to 0.5–0.7, while a broad particle size distribution of formulations has been shown with a PDI value > 0.7. The PDI values of 0.54 (±0.013) for SPPFQ and 0.79 (±0.022) for SPPFE indicate a difference in the uniformity of the nanocapsules, with the SPPFQ nanocapsules showing a more homogeneous distribution and the SPPFE nanocapsules showing a broader dispersion, which is probably related to the difference in the incorporated compounds. The zeta potential plots (**Figure 11 B** and **D**) show that both types of nanocapsules have a negative surface charge, which is related to the presence of FA in both types of nanocapsules. The lower negative zeta potential of the SPPFE nanocapsules may be due to the presence of plant compounds with cathodic electromagnetic properties (positive charge) next to FA, which partially neutralize the surface charge, which can affect the colloidal stability and interactions with biological systems. The intensity-based size distribution plots (**Figure 11 A** and **C**) show a monomodal peak for both types of nanocapsules, confirming the absence of large aggregates. However, the broader peak for SPPFE is consistent with its higher PDI, indicating greater size variability. The zeta potential distributions (**Figure 11 B** and **D**) are symmetric, indicating consistent surface charge characteristics across populations.

### Release pattern of essential oils or quercetin from nanocapsules

The release kinetics data in **Figure 12** show distinct pH-responsive profiles for SPPPFE (essential oil) and SPPFQ (quercetin) nanocapsules. Under physiological conditions (pH=7.4), both systems exhibit gradual release, reaching approximately 40% (SPPFE) and approximately 30% (SPPFQ) after 72 h, indicating that stable encapsulation is suitable for systemic circulation. At acidic pH (pH=5.5), the release rate increases dramatically, reaching approximately 80% (SPPFE) and approximately 70% (SPPFQ) within 24 h, which is attributed to polymer swelling due to protonation and increased drug solubility. The initial burst release (15–60 min) at pH=5.5 indicates surface-bound cargo release followed by sustained diffusion. The time-dependent curves highlight the faster release of SPPFE than of SPPFQ, which is probably due to the lower molecular weight of the essential oil. These results confirmed the biphasic release of the nanocapsules.

**Figure 12.**
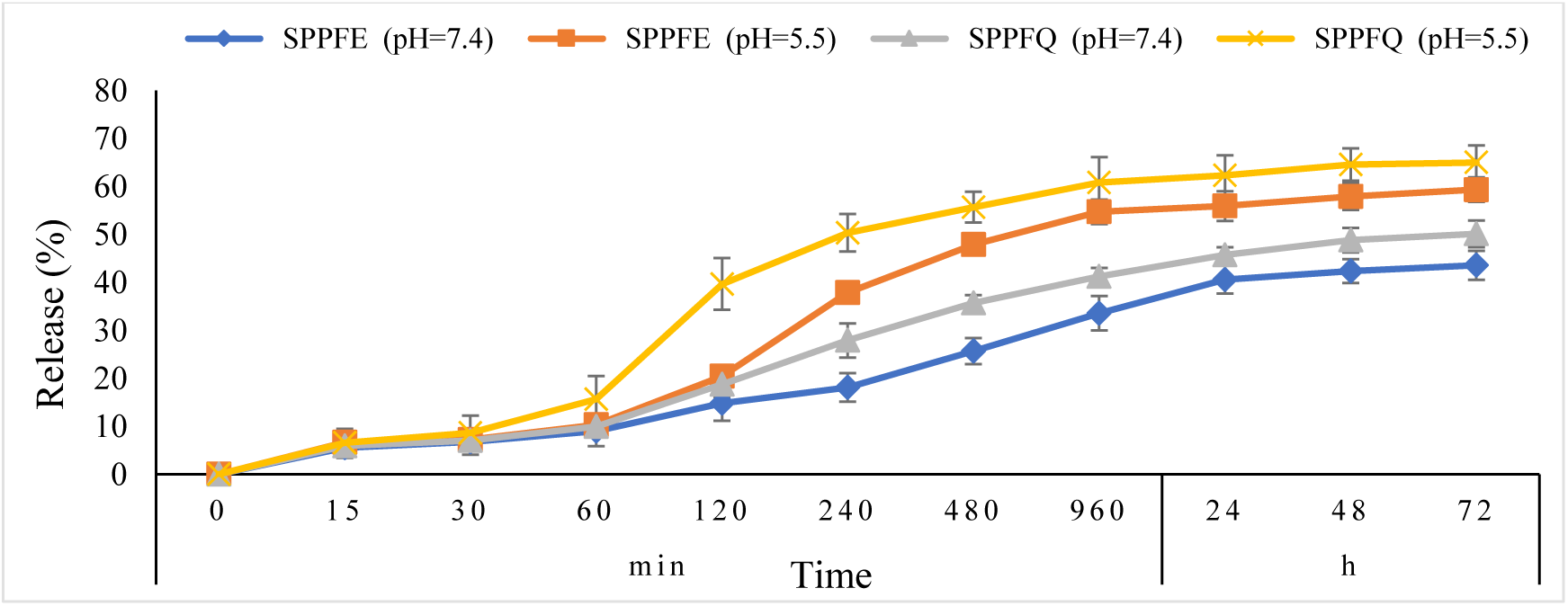
Essential oil or quercetin release patterns from SPPFE and SPPFQ nanocapsules at different pH values (pH=7.4 and pH=5.5).

### Anticancer properties

#### Cell viability

The cytotoxic effects of EO, Que and different nanocapsule formulations (SPPF, SPPFE and SPPFQ) at different concentrations on MCF-7 gastric cancer cells were evaluated (**Figure 13**). The results revealed that EO and Que (in free form) showed dose-dependent cytotoxicity, with a significant decrease in cell viability observed with increasing concentrations of each compound, reaching 23.04% and 11.03% viability at 100 μg/mL, respectively. In contrast, the SPPF nanoparticle formulation showed minimal cytotoxicity at all the tested doses, indicating the high biocompatibility of this group of nanoparticles. The encapsulated SPPPFE and SPPFQ formulations showed high toxicity profiles, with cell viability decreasing to 20.08% and 9.09% at 100 μg/mL, respectively. These findings suggest that nanoencapsulation can enhance the intrinsic cytotoxicity of quercetin and EO through targeted cellular binding. Statistical analysis via Duncan’s test confirmed significant differences between treatments (p<0.05), with letters indicating homogeneous groups. These results support the potential of nanoformulations in reducing the side effects of natural compounds. SPPF nanoparticles stand out as promising nontoxic candidates for further therapeutic development.

**Figure 13.**
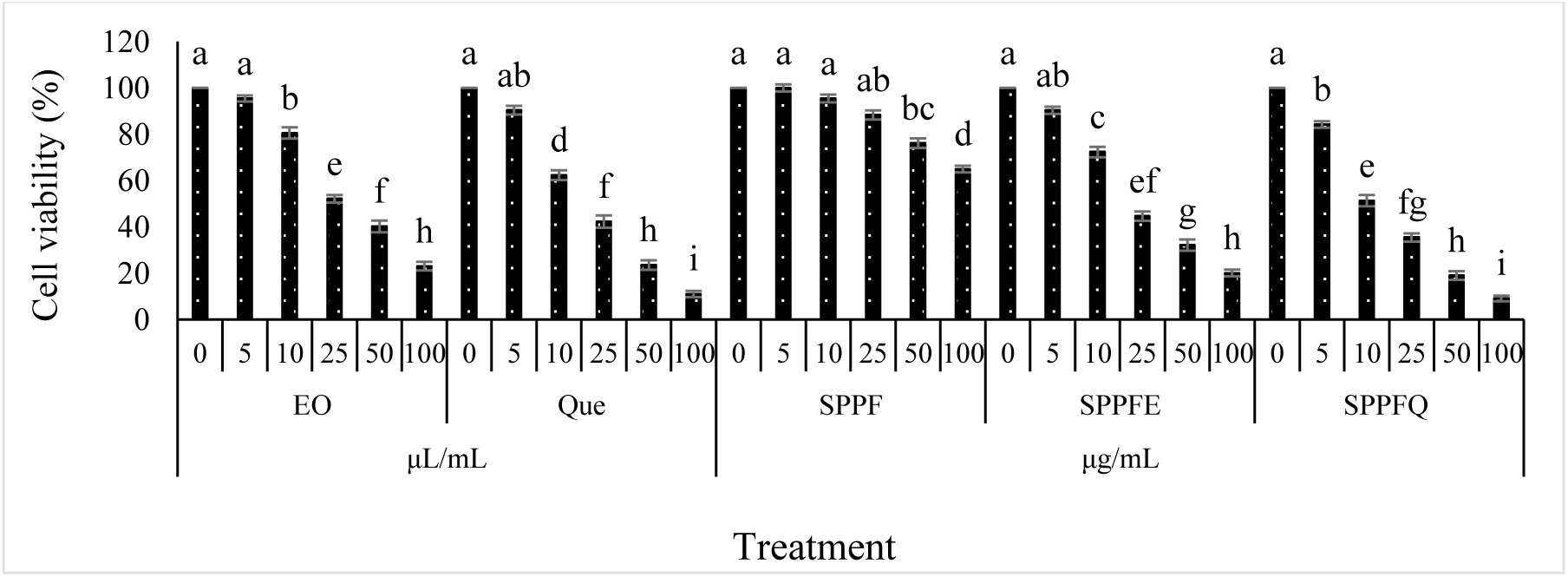
Toxicity of the EO (essential oil), Que (quercetin), SPPF, SPPFE, and SPPFQ nanocapsules to MCF-7 cells. *Lowercase letters (a, b, c) above the columns/graphs indicate the results of the statistical tests (Duncan, ANOVA). Common letters indicate no significant difference. Different letters indicate significant differences (p<0.05).

On the other hand, the results of this study (**Figure 13**) show that different nanocapsule formulations have different effects on MCF-7 cell viability. The results of the statistical tests revealed that the nanocapsule containing quercetin (SPPFQ) had the greatest cytotoxic effect, which was probably due to its antioxidant properties and the induction of apoptosis by this compound. In contrast, other treatment methods, such as pure essential oil and SPPF and SPPPFE nanocapsules, resulted in less toxicity. The data analysis revealed that the significant differences between the groups (p<0.05) could be due to differences in cell membrane permeability, controlled drug release or specific interactions between the active compounds and cancer cells. Additionally, a dose‒response relationship was observed for some of the treatments.

#### IC_50_

According to the IC_50_ results of EO, Que and nanocapsules (SPPF, SPPFE, and SPPFQ) on the MCF-7 cell line (**Figure 14**), SPPFQ had the lowest IC_50_ (IC_50_=11.21 μg/mL), whereas the free form of Que also had a lower IC_50_ (IC_50_=14.67 μL/mL) (**Figure 15 B** and **E**). In other words, quercetin, both free and nanocapsulated, had the strongest effects. In contrast, the EO had the weakest inhibitory effect (IC_50_=27.77 μL/mL), which was probably due to differences in its composition or bioavailability (**Figure 15 A**). The superior performance of SPPFQ over SPPFE (IC_50_=23.08 μg/mL) highlights the synergistic role of quercetin in nanocapsule formulations (**Figure 15 D** and **E**). These findings suggest potential applications in drug delivery systems, particularly for anticancer therapies.

**Figure 14.**
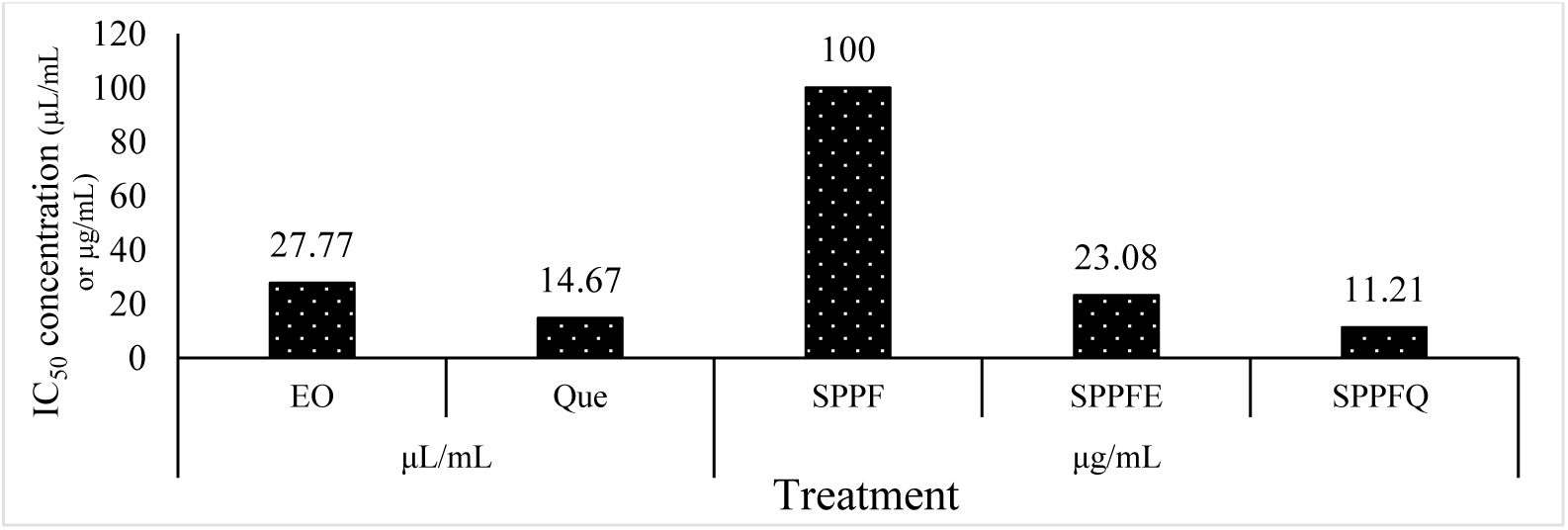
IC50 concentrations of the EO (essential oil), Que (quercetin), SPPF, SPPFE, and SPPFQ nanocapsules.

**Figure 15.**
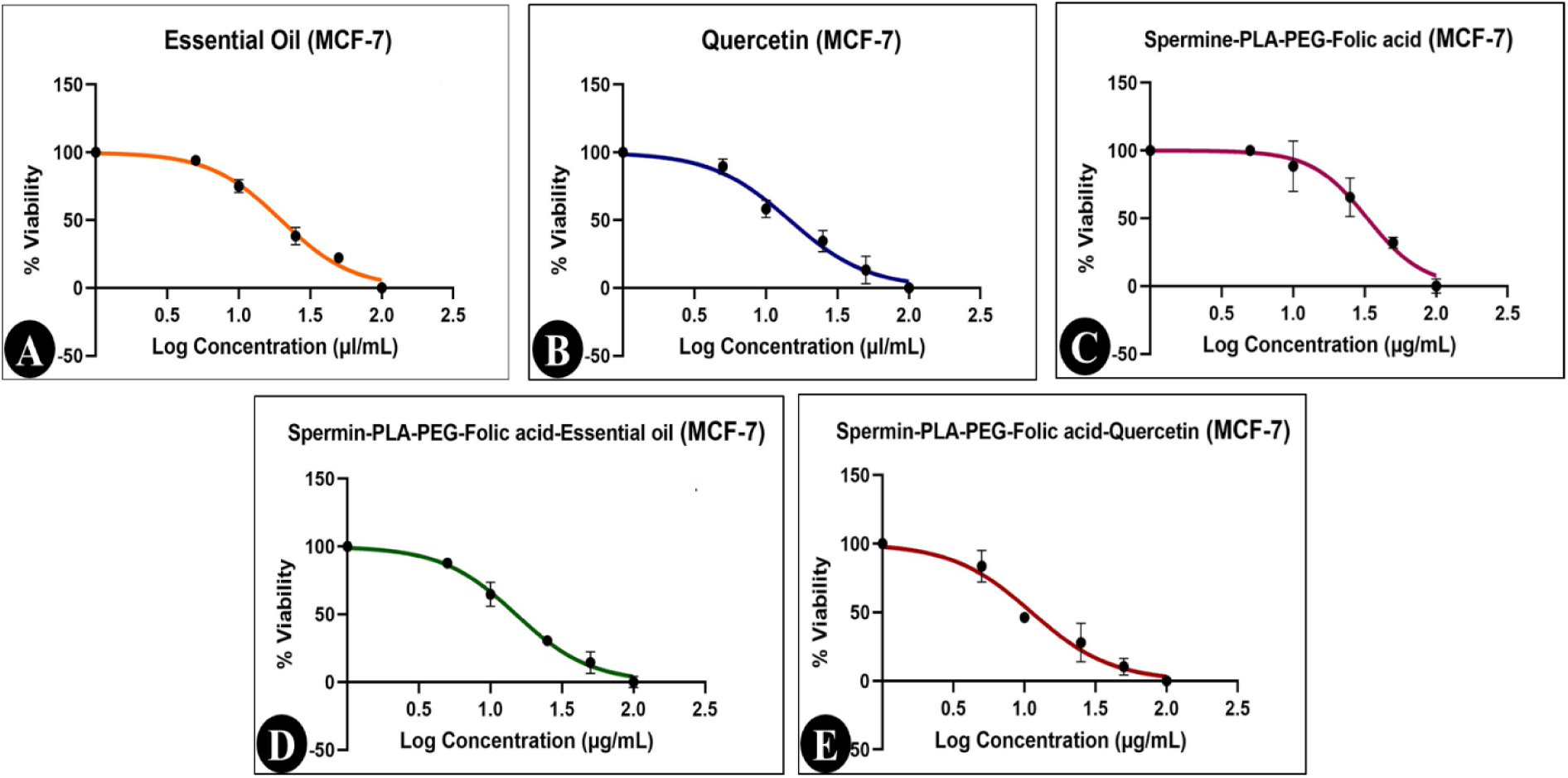
IC50 concentration chart of **A**) essential oil, **B**) quercetin, **C**) SPPF, **D**) SPPFE, and **E**) SPPFQ nanocapsules.

#### Apoptosis

According to the flow cytometry results (**Figures 16** and **17**), there was a significant difference between the different treatments (with IC_50_ values) in terms of the percentage of necrotic, preapoptotic, and postapoptotic cells. In terms of the percentage of necrotic cells, the highest percentage of necrotic cells (23.5%) was in the control sample (without treatment) (**Figure 17 A**), and the lowest percentage (12.2%) was in the cells treated with SPPFQ nanocapsules (**Figure 17 E**). With respect to the percentage of pre– and postapoptotic cells, a statistically significant difference was observed between the treatments in the MCF-7 cell line treated with the values obtained from the IC_50_ test. Thus, the highest percentage of pre– and postapoptotic cells was detected in the MCF-7 cells treated with SPPFQ nanocapsules (10.78 and 20.07%, respectively) (**Figures 16** and **17 E**).

**Figure 16.**
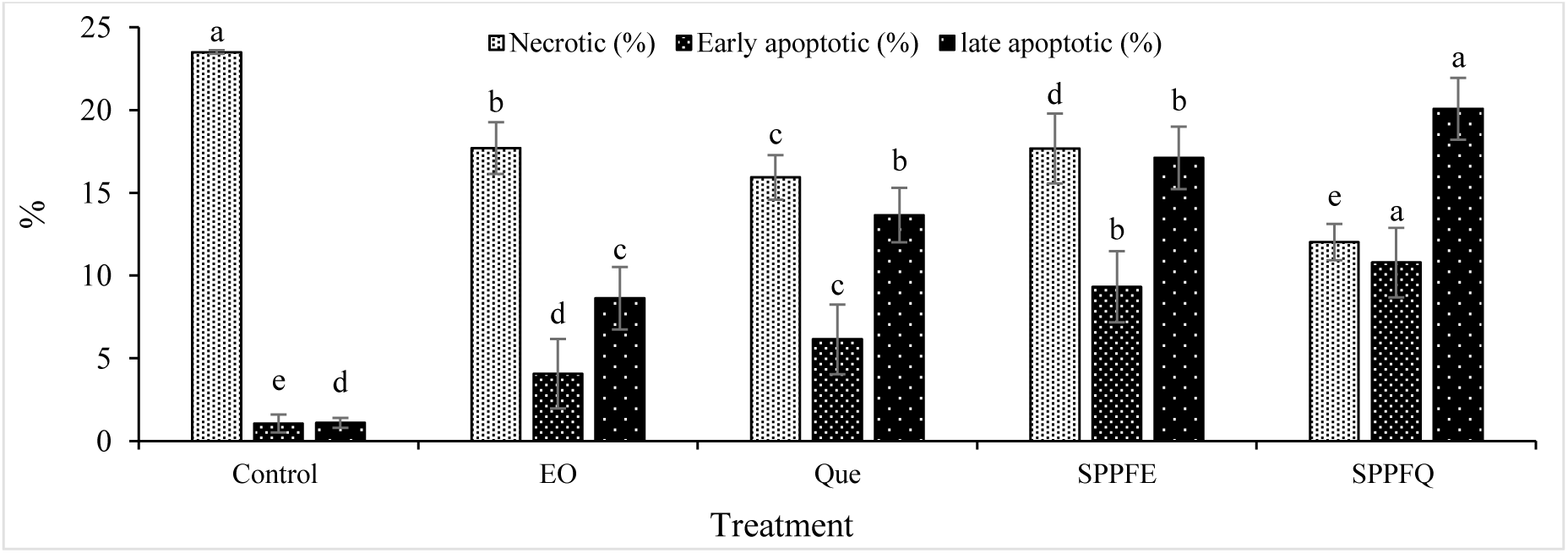
Effects of EO (the essential oil of *P. oleracea*), Que (quercetin), SPPF, SPPFE, and SPPFQ nanocapsules on the percentage of necrotic cells, the preapoptotic stage, and the postapoptotic stage of the MCF-7 cell line.

**Figure 17.**
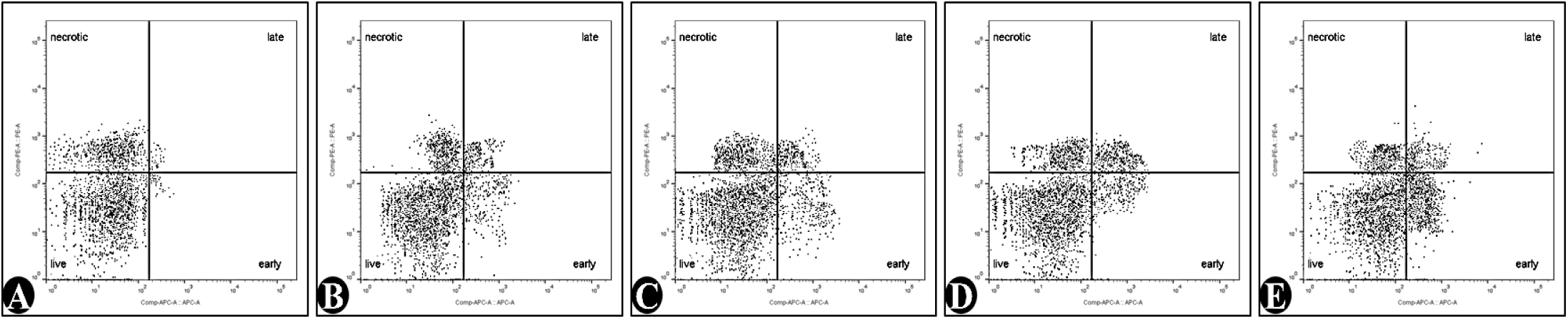
Flow cytometry results: MCF-7 cell line treated; **A)** control treatment, **B)** essential oil of *P. oleracea* (EO), **C)** pure quercetin (Que), **D)** SPPFE, and **E)** SPPFQ

As shown in Figure 16, the use of Que and *P. oleracea* EO in free and nonencapsulated forms led to a significant increase in the percentage of necrotic cells compared with that of cells in the preapoptotic and postapoptotic stages. This pattern of cell death is mainly nontargeted and is caused by mechanisms of general toxicity and nonspecific destruction (**Figures 16** and **17 B, C**).

In contrast, the formulation of these compounds in the form of SPPFE and SPPFQ nanocapsules (containing EO and Que, respectively) significantly changed the pattern of cell death. The results show that encapsulation, especially in samples containing folic acid, leads to a significant increase in the population of preapoptotic and postapoptotic cells compared with necrotic cells. This change in pattern indicates the activation of targeted apoptotic pathways and a decrease in random cell death (**Figure 17 D** and **E**).

In particular, the SPPFQ nanocapsules (containing quercetin) showed better apoptosis induction performance than did SPPFE (containing essential oil), which is likely due to the intrinsic ability of quercetin to modulate apoptotic signals. These findings emphasize the advantages of nanocarrier technology in enhancing the efficacy and specificity of natural anticancer compounds. The use of these smart drug delivery systems could be a promising strategy to reduce side effects and increase the efficacy of herbal-based therapies.

## Discussion

Copolymer nanoparticles are an innovative approach for delivering drugs and genetic materials to cells that offer several advantages for therapeutic applications. These nanoparticles are highly compatible with biological systems, and their surface properties can be precisely tuned to optimize their performance (Xia et al., 2021). A key feature of the specific polymers used in these nanoparticles is their positive charge, which allows them to effectively bind to drug compounds and genetic materials such as DNA, a molecule that has a negative charge (Mohajeri, Dashti, et al., 2025). This interaction promotes efficient entry into cells through a natural uptake process known as endocytosis (Banushi et al., 2023). Compared with conventional cancer treatments such as chemotherapy, which often cause severe side effects, gene or drug delivery via these polymeric systems offers a milder alternative (Mansuryar et al., 2025). This technology has significant potential to enhance treatment outcomes and improve the overall well-being of cancer patients by minimizing harmful side effects (de Almeida et al., 2021).

Natural polyamines such as polyspermine have emerged as promising candidates for drug delivery because of their biocompatibility, drug affinity, and structural advantages. The polycationic nature of polyspermine allows for efficient encapsulation of therapeutic drugs, whereas their pH-responsive behavior allows for controlled release into tumor microenvironments (Mohajeri, Fathi Erdi, & Yaghoubi, 2025). To enhance tumor targeting, we incorporated folic acid—a known ligand that binds to folate receptors that are overexpressed in cancer cells—into spermine-PLA-PEG copolymer nanoparticles.

Before evaluating the anticancer efficacy, we characterized purslane essential oils via GC‒MS and identified key bioactive compounds with significant antioxidant and antimicrobial properties. These nanocapsules were then engineered for targeted drug delivery of purslane oil and quercetin to MCF-7 breast cancer cells, utilizing receptor-mediated uptake and pH-triggered release to maximize therapeutic effects while minimizing off-target toxicity.

This approach is in line with recent advances in ligand-functionalized nanomedicine and demonstrates how natural polymers and tumor-localized moieties can work together to improve drug delivery precision.

The bioactive properties of *P. oleracea* essential oils were systematically evaluated through comprehensive antimicrobial and antioxidant assays. Our disk diffusion results (**Figure 4**) demonstrated concentration-dependent growth inhibition against both *S. aureus* and *E. coli*, with significantly greater efficacy against gram-positive pathogens (p<0.01). These findings align with previous reports on the increased susceptibility of gram-positive bacteria to plant essential oils due to their simpler cell wall structure (Andrade-Ochoa et al., 2021).

The MIC/MBC results (**Figure 5**) revealed notable differences in bacterial sensitivity, with *S. aureus* showing twice the susceptibility of *E. coli*. This pattern corresponds with recent studies on essential oil mechanisms of action, where the outer membrane of gram-negative bacteria provides additional resistance (Ike et al., 2025). The observed MIC values (12.5–25 μL/mL) compare favorably with those reported for other medicinal plant essential oils in the contemporary literature (Zhang et al., 2025).

Our DPPH radical scavenging assessment (**Figure 6**) revealed dose-dependent antioxidant activity, reaching 27.61% inhibition at 100 μL/mL. While this activity appears to be moderate compared with that of synthetic antioxidants, it remains significant considering the complex mixture of phytochemicals identified through GC‒ MS analysis (**Table 1**). These results support the growing body of evidence that *P. oleracea* contains bioactive compounds with potential therapeutic applications (Kumar et al., 2022).

Structural and compositional characterization of the synthesized polyspermine-PLA-PEG/folic acid copolymer was performed via multiple analytical techniques. The ^1^H-NMR, FTIR, TGA, and DTG analyses provided conclusive evidence of successful polymer formation and functional group incorporation (**Figures 7, 8, and 9**).

Previous studies have reported similar synthetic approaches for folate-conjugated copolymers. Wadhawan et al. (2022) demonstrated the successful attachment of folic acid to PLA-PEG-polyspermine copolymers (Wadhawan et al., 2022), with characteristic aromatic proton resonances appearing between 6-8 ppm in the ¹H NMR spectrum, a finding that aligns with our experimental results (**Figure 7**). Complementary evidence comes from recent work by Ramalho et al. (2024), who developed folate-functionalized PLGA nanoparticles. Their FTIR analysis revealed distinctive absorption bands at 1750 cm^-1^ (carbonyl stretch) and 1650 cm^-1^ (aromatic vibrations), mirroring the spectral features observed in our PSPF copolymer system (Ramalho et al., 2024). These collective findings from independent research groups validate our synthetic methodology and confirm the expected chemical structure of the developed copolymer (**Figure 8**).

Recent advances in nanocarrier design have highlighted the critical importance of thermal stability in ensuring pharmaceutical product integrity. Our thermal characterization studies, which are consistent with the findings of Javed et al. (2022), demonstrate that copolymer nanostructures exhibit distinct degradation patterns that correlate with their structural composition (Javed et al., 2022). The thermal profiles observed in our work (**Figure 9**) align with established literature reports on similar polymeric systems (Lei et al., 2021), particularly regarding the characteristic multiphase degradation of multicomponent copolymers (Lei et al., 2021). The enhanced thermal resilience observed in our nanocapsules mirrors the results reported by Jiang et al. (2024) for analogous copolymer architectures (Jiang et al., 2024). As shown in **Figure 9 C**, the improved stability profile compared with that of individual components confirms successful copolymer formation, a phenomenon well documented in recent studies of polyamine-integrated systems (Holbert et al., 2022). These findings support the growing body of evidence that strategic copolymer design can significantly enhance thermal properties (Constantinou et al., 2023). Our DTG analysis (**Figure 9 D-F**) revealed degradation kinetics consistent with those reported in comprehensive reviews of polymeric carrier systems (Visan et al., 2021). The complex multistep degradation profile particularly matches observations by Tune et al. (2022) for similar nanostructures, suggesting comparable molecular interactions within the copolymer matrix (Tune et al., 2022).

These thermal characteristics, as demonstrated in our results (**Figure 9**), meet the stability benchmarks proposed by Domingues et al. (2023) for pharmaceutical applications (Domingues et al., 2023). The structural integrity observed at biologically relevant temperatures suggests promising shelf-life stability, a crucial factor emphasized in current good manufacturing practice guidelines (Domingues et al., 2023).

The shape and morphology of nanoparticles, as key factors in the targeted gene transfer process, are highly important and significantly affect the efficiency of these delivery systems.

TEM analysis revealed critical insights into the structural properties of the synthesized SPPFE and SPPFQ nanocapsules (**Figure 10**). Both formulations exhibited well-defined spherical morphologies, characteristic of polymeric nanoparticles fabricated through emulsion-based techniques. The observed particle diameters fell within the nanoscale range (200 nm scale bar), demonstrating successful control over the particle size distribution during synthesis.

Notably, the TEM micrographs revealed distinct dark cores within the nanoparticles, providing visual confirmation of essential oil encapsulation within the polymer matrix. Comparative analysis between the two formulations revealed subtle morphological differences: SPPFQ nanocapsules (**Figure 10 B**) exhibited more uniform shape definitions than did their SPPFE counterparts (**Figure 10 A**). This enhanced structural homogeneity in SPPFQ may be attributed to the incorporation of quercetin, whose polyhydroxylated structure potentially influences polymer chain packing during nanoparticle formation.

The absence of particle aggregation in both systems underscores the effectiveness of PEG as a stabilizing agent, which is consistent with previous reports by Zhou et al. (2023) on PEGylated nanocarriers (Zhou et al., 2023). Our findings align with the work of Noruzpour et al. (2024), who similarly reported that compared with essential oil-loaded systems, flavonoid-containing nanoparticles (quercetin derivatives) demonstrated improved morphological regularity (Noruzpuor et al., 2024). The uniform electron density distribution throughout the nanoparticles suggests homogeneous dispersion of the encapsulated actives, mirroring the results reported by Mohajeri et al. (2025) for analogous polymer-based delivery systems (Mohajeri, Yaghoubi, et al., 2025). The well-dispersed nature of our nanocapsules, combined with their demonstrated loading capacity, make them promising candidates for targeted delivery applications.

Recent advances in nanocarrier technology have demonstrated the critical importance of physicochemical properties in determining the biological behavior of drug delivery systems. The current findings align with established principles in nanomedicine, where particle size and surface charge significantly influence biodistribution and cellular uptake mechanisms (Amani et al., 2021). The observed size range of 115–237 nm in our systems corresponds (**Table 2** and **Figure 11 A** and **C**) well with the optimal diameter window (50–300 nm) for enhanced tumor accumulation via the EPR effect, as extensively documented in cancer nanotherapeutics (Sh et al., 2024).

The differential behavior between flavonoid-loaded and essential oil-encapsulated systems reflects a broader trend in nanocarrier design. As noted by Noruzpour et al. (2025), single bioactive compounds typically yield more uniform nanostructures than complex plant extracts do because of their defined molecular characteristics. This phenomenon has been particularly well characterized in polymeric nanosystems, where the molecular weight and hydrophobicity of encapsulated agents directly impact nanoparticle morphology and stability (Noruzpour et al., 2025).

The surface charge characteristics observed in our study (−6.5 to –8.2 mV) (**Table 2** and **Figure 11 B** and **D**) warrant special consideration in light of recent developments in stealth nanoparticle technology. While strongly negative or positive charges often promote rapid clearance, moderate surface potentials in this range are associated with favorable circulation times. This is particularly relevant given the increasing focus on folic acid-functionalized systems for targeted delivery, where surface properties must balance stealth characteristics and receptor recognition (Bourang et al., 2025).

The differences in polydispersity between formulations highlight their importance in nanomedicine translation. As emphasized by Xie et al. (2022), batch-to-batch consistency represents a critical quality attribute for clinical translation, with PDI values below 0.3 being ideal for injectable formulations (Xie et al., 2022). While our systems show higher values, this is consistent with phytochemical-loaded systems where natural product variability often challenges perfect monodispersity (Bourang, Jahanbakhsh Godehkahriz, et al., 2024).

These findings contribute to the increasing understanding of structure‒function relationships in nanocarrier design, particularly for plant-derived therapeutics. The demonstrated ability to encapsulate both single flavonoids and complex essential oils within similar polymeric architectures provides valuable insights for developing versatile delivery platforms. Future directions should explore how these physicochemical differences translate to in vivo performance, particularly in terms of pharmacokinetics and target site accumulation, an area where the current literature shows significant gaps (Lin et al., 2022).

The pH-dependent release behavior observed in our study (**Figure 12**) provides compelling evidence for the stimuli-responsive properties of these nanocapsule formulations. Our findings align with established principles in nanocarrier design, where pH-triggered drug release has emerged as a key strategy for targeted delivery (Singh & Nayak, 2023). The differential release patterns between SPPFE and SPPFQ nanocapsules under varying pH conditions mirror observations reported in recent studies (Bourang, Asadian, et al., 2024).

The sustained release profile at physiological pH (pH=7.4) demonstrated excellent formulation stability, which was consistent with findings from Haseli et al. (2022) on similar copolymer systems (Haseli et al., 2022). Our results showing approximately 40% (SPPFE) and 30% (SPPFQ) release after 72 hours correlate well with the optimal release kinetics for systemic circulation proposed by Lakshani et al. (2023) (Lakshani et al., 2023). This controlled release behavior is particularly advantageous for maintaining therapeutic drug levels while minimizing side effects (Wang et al., 2021).

The accelerated release observed at acidic pH (pH=5.5) supports the growing body of evidence for pH-responsive polymer behavior. Our data, which revealed 80% (SPPFE) and 70% (SPPFQ) release within 24 h, confirm the protonation-induced swelling mechanism first described by Negrón et al. (2025) (Negrón et al., 2025). The biphasic release pattern we observed, featuring an initial burst followed by sustained release, has been widely reported as characteristic of well-designed nanocarrier systems (Altuwaijri et al., 2025). These findings have significant implications for cancer therapy applications, where the acidic tumor microenvironment can be exploited for targeted drug release. Our results corroborate recent clinical observations that pH-responsive systems can enhance therapeutic efficacy while reducing systemic toxicity (Verkhovskii et al., 2023).

The evaluation of the cytotoxic effects of our nanocapsule formulations against MCF-7 breast cancer cells reveals important structure‒activity relationships that align with current trends in nanomedicine development. The observed dose-dependent cytotoxicity of free essential oil (EO) and quercetin (**Figure 13**) corresponds with previous reports on plant-derived anticancer agents (Bourang et al., 2025). However, the significantly enhanced activity of the encapsulated forms (SPPFE and SPPFQ) supports the growing body of evidence that nanoformulations can potentiate the efficacy of bioactive compounds.

Our IC_50_ results (**Figures 14** and **15**) demonstrate the remarkable potency of quercetin-loaded nanocapsules (11.21 μg/mL), which is consistent with recent findings on flavonoid nanoformulations (Jasim et al., 2022). The superior performance of SPPFQ over free quercetin (14.67 μg/mL) suggests improved cellular uptake, as reported in similar delivery systems (Wang et al., 2023).

Flow cytometry analysis (**Figures 16** and **17**) provided compelling evidence that apoptosis induction is the primary mechanism of action, particularly for SPPFQ nanocapsules. These findings corroborate recent studies demonstrating that nanoencapsulation can shift cell death mechanisms from necrosis to apoptosis (Zhang et al., 2023). The observed apoptotic populations (10.78% early, 20.07% late apoptosis) significantly exceeded the values reported for free compounds in similar systems (Tomou et al., 2023). The differential response between the SPPFE and SPPFQ formulations highlights the importance of payload characteristics in nanocarrier design. While both systems showed improved efficacy over free compounds, the superior performance of quercetin-loaded nanocapsules aligns with their well-documented proapoptotic signaling pathways. This observation supports the emerging paradigm of combining natural compounds with advanced delivery technologies to maximize therapeutic potential.

## Conclusion

This study successfully developed spermine-PLA-PEG-FA copolymer nanoparticles functionalized with folic acid for targeted delivery of *P. oleracea* essential oil and quercetin to cancer cells. Our approach combines the inherent biocompatibility of natural polyamines with tumor-targeting ligands to increase therapeutic precision while minimizing systemic toxicity. Structural characterization confirmed successful nanoparticle synthesis through multiple analytical techniques, with thermal analysis demonstrating enhanced stability of the copolymer system compared with that of individual components. The nanocapsules exhibited favorable physicochemical properties, including a nanoscale size (115-237 nm), moderate surface charge (−6.5-8.2 mV), and pH-responsive release kinetics—controlled release at physiological pH (pH=7.4) and accelerated drug release under acidic tumor conditions (pH=5.5). Biological evaluations revealed that the anticancer activity of quercetin-loaded nanoparticles (IC_50_ = 11.21 μg/mL) was superior to that of free compounds, and flow cytometry confirmed that apoptosis was the primary mechanism of cell death (20.07% late apoptosis). The essential oil components demonstrated significant antimicrobial effects, particularly against gram-positive bacteria (*E. coli*), along with notable antioxidant activity. These findings collectively validate our nanocarrier design as a promising platform for targeted cancer therapy, offering improved payload delivery, enhanced therapeutic efficacy, and reduced off-target effects compared with conventional treatments. This study provides a foundation for the further development of plant-based nanomedicines, with potential applications extending beyond oncology to other therapeutic areas requiring targeted delivery systems.

## Author contributions

MN and SB conducted the experiments, analyzed the data, and wrote the original draft. RAZ conceived and designed the research, administered and supervised the project, and reviewed and edited the manuscript. HY collaborated in the implementation of the HPLC analysis and flow cytometry. SB and MN contributed new analytical tools and revised the manuscript. All the authors read and approved the manuscript.

## Ethics approval

This article does not contain any studies with human participants or animals performed by any of the authors. The MCF-7 cell line and the bacterial strains *Staphylococcus aureus* and *Escherichia coli* used in this study were obtained from the American Type Culture Collection (ATCC), and their use does not require ethical approval.

## Consent to participate

Not applicable.

## Consent for publication

Not applicable.

## Competing interests

The authors declare that they have no competing interests.

## Acknowledgment

This work was supported by the Office of the Vice-Chancellor for Research and Technology of the University of Mohaghegh Ardabili.

## Funding

This research did not receive any specific grant from funding agencies in the public, commercial, or not-for-profit sectors.

## Data availability

The authors declare that the data supporting the findings of this study are available within the paper. Should any raw data files be needed in another format, they are available from the corresponding author upon reasonable request.

## Abbreviations

DPPH: 2,2-diphenyl-1-picrylhydrazyl
MTT: 3-(4,5-dimethylthiazol-2-yl)-2,5-diphenyltetrazolium bromide
DTG: Differential thermogravimetric analysis
DMSO: Dimethyl sulfoxide
DLS: Dynamic light scattering
EO: Essential oil
FBS: Fetal bovine serum
FTIR: Fourier transform infrared spectroscopy
GC–MS: Gas chromatography–mass spectrometry
MBC: Minimum bactericidal concentration
MIC: Minimum inhibitory concentration
PDI: Polydispersity Index
PVA: Polyvinyl alcohol
1H-NMR: Proton nuclear magnetic resonance spectroscopy
Que: Quercetin
RES: Reticuloendothelial system
SPPFE: Spermine-PLA-PEG-FA/Essential Oil
SPPFQ: Spermine-PLA-PEG-FA/Quercetin
SPPF: Spermine-PLA-PEG-FA
Spermin-PLA-PEG-FA: Spermine-polylactic acid-polyethylene glycol
TGA: Thermogravimetric analysis
TEM: Transmission electron microscopy

## Notes

### Competing Interest Statement

The authors have declared no competing interest.

## References

1. Ahmadi-Nouraldinvand F, et al. (2024). Preparation and characterization of multi-target nanoparticles for co-drug delivery. Medicine in Drug Discovery, 21, 100177.

2. Al Kamaly O, et al. (2024). Phytochemical Composition and Cytotoxic Activity of Senecio asirensis Hexane Fraction Using In Vitro and In Silico Approaches. Natural Product Communications, 19(4), 1934578X241246418.

3. Alonso-Castro A J, et al. (2022). Myristic acid reduces skin inflammation and nociception. Journal of Food Biochemistry, 46(1), e14013.

4. Alshawwa S Z, et al. (2022). Nanocarrier drug delivery systems: characterization, limitations, future perspectives and implementation of artificial intelligence. Pharmaceutics, 14(4), 883.

5. Altuwaijri N, et al. (2025). Assessing the Antibacterial Potential and Biofilm Inhibition Capability of Atorvastatin-Loaded Nanostructured Lipid Carriers via Crystal Violet Assay. Pharmaceuticals, 18(3), 417.

6. Amani A, et al. (2019). Multifunctional magnetic nanoparticles for controlled release of anticancer drug, breast cancer cell targeting, MRI/fluorescence imaging, and anticancer drug delivery. Journal of Drug Delivery Science and Technology, 49, 534–546.

7. Amani A, et al. (2021). Design and invitro characterization of green synthesized magnetic nanoparticles conjugated with multitargeted poly lactic acid copolymers for co-delivery of siRNA and paclitaxel. European Journal of Pharmaceutical Sciences, 167, 106007.

8. Andrade-Ochoa S, et al. (2021). Differential antimicrobial effect of essential oils and their main components: Insights based on the cell membrane and external structure. Membranes, 11(6), 405.

9. Arnold M, et al. (2022). Current and future burden of breast cancer: Global statistics for 2020 and 2040. The Breast, 66, 15–23.

10. Asami E, et al. (2023). Anti-inflammatory activity of 2-methoxy-4-vinylphenol involves inhibition of lipopolysaccharide-induced inducible nitric oxidase synthase by heme oxygenase-1. Immunopharmacology and Immunotoxicology, 45(5), 589–596.

11. Asghari Zakaria R, et al. (2024). Investigating the anticancer properties of the essential oil and aqueous extract of Moringa oleifera and its biosynthesized metal nanoparticles on MCF-7 and BT-549 cell lines. Iranian Journal of Breast Diseases, 17(1), 59–83.

12. Banushi B, et al. (2023). Endocytosis in cancer and cancer therapy. Nature Reviews Cancer, 23(7), 450–473.

13. Barreiro-Costa O, et al. (2021). Synthesis and evaluation of biological activities of bis (spiropyrazolone) cyclopropanes: A potential application against leishmaniasis. Molecules, 26(16), 4960.

14. Barros A d L, et al. (2022). Cytotoxicity and lipase inhibition of essential oils from amazon annonaceae species.

15. Bhandari S, et al. (2011). Determining the limits and confounders for the 2-pentyl furan breath test by gas chromatography/mass spectrometry. Journal of Chromatography B, 879(26), 2815–2820.

16. Bourang S, et al. (2024). PLA-HA/Fe3O4 magnetic nanoparticles loaded with curcumin: physicochemical characterization and toxicity evaluation in HCT116 colorectal cancer cells. Discover Applied Sciences, 6(4), 186.

17. Bourang S, et al. (2024). Green synthesis of iron oxide, copper, zinc oxide and silver nanoparticles from aqueous extract of F. vulgare and evaluation of their structural and antimicrobial properties. Agricultural Biotechnology Journal, 16(3), 60–88.

18. Bourang S, et al. (2025). Anticancer properties of copolymer nanoparticles loaded with Foeniculum vulgare derivatives in Hs578T and SUM159 cancer cell lines. Cancer Nanotechnology, 16(1), 1–28.

19. Bourang S, et al. (2024). Synthesis and in vitro characterization of PCL-PEG-HA/FeCo magnetic nanoparticles encapsulating curcumin and 5-FU. Nanomedicine Journal, 11(2).

20. Bourang S, et al. (2024). Application of nanoparticles in breast cancer treatment: a systematic review. Naunyn-Schmiedeberg’s Archives of Pharmacology, 1–47.

21. Chiriac A P, et al. (2021). Polymeric carriers designed for encapsulation of essential oils with biological activity. Pharmaceutics, 13(5), 631.

22. Cimino C, et al. (2021). Essential oils: Pharmaceutical applications and encapsulation strategies into lipid-based delivery systems. Pharmaceutics, 13(3), 327.

23. Constante C K, et al. (2022). Adaptation of the methyl thiazole tetrazolium (MTT) reduction assay to measure cell viability in Vibrio spp. Aquaculture, 560, 738568.

24. Constantinou A P, et al. (2023). Thermoresponsive block copolymers of increasing architecture complexity: a review on structure–property relationships. Polymer Chemistry, 14(3), 223–247.

25. Cordeiro M E R, et al. (2023). Effects of α-Humulene and its Nanoparticles in Experimental Model of Alzheimer’s Disease: Behavioral Analysis and Anti-Inflammatory Activity. Journal of Advances in Medicine and Medical Research, 35(24), 1–13.

26. Cui R, et al. (2022). GC-MS analysis and anti-inflammatory activity of low polarity parts from 3 species of Sabia genus. China Pharmacy, 446–451.

27. de Almeida M S, et al. (2021). Understanding nanoparticle endocytosis to improve targeting strategies in nanomedicine. Chemical society reviews, 50(9), 5397–5434.

28. Domingues C, et al. (2023). Pediatric drug development: reviewing challenges and opportunities by tracking innovative therapies. Pharmaceutics, 15(10), 2431.

29. Donaldson J R, et al. (2005). Assessment of antimicrobial activity of fourteen essential oils when using dilution and diffusion methods. Pharmaceutical biology, 43(8), 687–695.

30. Duanis-Assaf D, et al. (2023). Phenyl tetramethyl cyclopropane carboxamide class: new broad-spectrum postharvest fungicides. Food Control, 154, 110041.

31. Fernandes Melo Reis R C, et al. (2024). From clove oil to bioactive agents: synthetic routes, antimicrobial and antiparasitic activities of eugenol derivatives. Future Medicinal Chemistry, 1–20.

32. Fouda A, et al. (2022). Antimicrobial, antiviral, and in-vitro cytotoxicity and mosquitocidal activities of Portulaca oleracea-based green synthesis of selenium nanoparticles. Journal of Functional Biomaterials, 13(3), 157.

33. Freitas M R C. (2023). Internship Reports and Monograph entitled “Isoeugenol, a skin allergen with therapeutic potential in Diabetes mellitus*.”*

34. Giaquinto A N, et al. (2022). Breast cancer statistics, 2022. CA: a cancer journal for clinicians, 72(6), 524–541.

35. Goswami L, et al. (2022). Design and synthesis of eugenol/isoeugenol glycoconjugates and other analogues as antifungal agents against Aspergillus fumigatus. RSC Medicinal Chemistry, 13(8), 955–962.

36. Haseli S, et al. (2022). A novel pH-responsive nanoniosomal emulsion for sustained release of curcumin from a chitosan-based nanocarrier: emphasis on the concurrent improvement of loading, sustained release, and apoptosis induction. Biotechnology Progress, 38(5), e3280.

37. Holbert C E, et al. (2022). Polyamines in cancer: integrating organismal metabolism and antitumour immunity. Nature Reviews Cancer, 22(8), 467–480.

38. Hong L, et al. (2023). Nanoparticle-based drug delivery systems targeting cancer cell surfaces. RSC advances, 13(31), 21365–21382.

39. Ike K A, et al. (2025). Evaluating the Effect of an Essential Oil Blend on the Growth and Fitness of Gram-Positive and Gram-Negative Bacteria. Biology, 14(4), 437.

40. Jahazi S, & Akbari, H. (2020). Preparation and characterization of doxorubicin loaded Fe3O4-PEG nanoparticles on AGS and MCF-7 cancer cells. Modares Journal of Biotechnology, 11(2), 167–175.

41. Janani K, et al. (2019). Chemical constituent, minimal inhibitory concentration, and antimicrobial efficiency of essential oil from oreganum vulgare against Enterococcus faecalis: An: in vitro: study. Journal of Conservative Dentistry and Endodontics, 22(6), 538–543.

42. Jang H-I, et al. (2020). Antibacterial and antibiofilm effects of α-humulene against Bacteroides fragilis. Canadian Journal of Microbiology, 66(6), 389–399.

43. Jasim A J, et al. (2022). Preliminary trials of the gold nanoparticles conjugated chrysin: An assessment of anti-oxidant, anti-microbial, and in vitro cytotoxic activities of a nanoformulated flavonoid. Nanotechnology reviews, 11(1), 2726–2741.

44. Javed S, et al. (2022). Nanostructured lipid carrier system: A compendium of their formulation development approaches, optimization strategies by quality by design, and recent applications in drug delivery. Nanotechnology reviews, 11(1), 1744–1777.

45. Javid S, et al. (2020). Semisynthesis of myristic acid derivatives and their biological activities: A critical insight. Journal of Biologically Active Products from Nature, 10(6), 455–472.

46. Jiang B, et al. (2024). Nanoparticle-Empowered Core–Shell Microcapsules: From Architecture Design to Fabrication and Functions. Small, 20(33), 2311897.

47. Karlinsey J E, et al. (2022). Cyclopropane fatty acids are important for Salmonella enterica serovar Typhimurium virulence. Infection and Immunity, 90(1), e00479–00421.

48. Keser F, et al. (2021). In vitro biological activities and phytochemical contents of Portulaca oleracea L.(Purslane). Journal of Physical Chemistry and Functional Materials, 4(1), 1–7.

49. Khammassi M, et al. (2024). Secondary metabolites of Santolina africana: chemical profiles and assessment of biological activities. International Journal of Secondary Metabolite, 11(3), 472–485.

50. Kita K, & Dittrich, C. (2011). Drug delivery vehicles with improved encapsulation efficiency: taking advantage of specific drug–carrier interactions. Expert opinion on drug delivery, 8(3), 329–342.

51. Kumar A, et al. (2022). A review on bioactive phytochemicals and ethnopharmacological potential of purslane (Portulaca oleracea L.). Heliyon, 8(1).

52. Lakshani N, et al. (2023). Release kinetic models and release mechanisms of controlled-release and slow-release fertilizers. ACS Agricultural Science & Technology, 3(11), 939–956.

53. Lei C, et al. (2021). Controlled vertically aligned structures in polymer composites: natural inspiration, structural processing, and functional application. Advanced Materials, 33(49), 2103495.

54. Lin Y H, & Chen, C-Y. (2020). Folate-targeted curcumin-encapsulated micellar nanosystem for chemotherapy and curcumin-mediated photodynamic therapy. Polymers, 12(10), 2280.

55. Lin Z, et al. (2022). Integration of in vitro and in vivo models to predict cellular and tissue dosimetry of nanomaterials using physiologically based pharmacokinetic modeling. ACS nano, 16(12), 19722–19754.

56. Manhas A, & Kaul, V. (2024). Metabolomics of Above and Below Ground Parts of Commelina benghalensis an Amphicarpic Weed Using GC–MS. National Academy Science Letters, 47(2), 187–193.

57. Mansuryar A, et al. (2025). The effect of Fe3O4 biosynthesized through the green synthesis of Silybum marianum and HA in the targeted delivery of 5-Fluorouracil to HCT116 cell line. DARU Journal of Pharmaceutical Sciences, 33(2), 27. 10.1007/s40199-025-00568-9

58. Martín-Sabroso C, et al. (2021). Active targeted nanoformulations via folate receptors: State of the art and future perspectives. Pharmaceutics, 14(1), 14.

59. Mohajeri S, et al. (2025). Design and preparation of PLA-chitosan-PEG-glucose copolymer for combined delivery of Paclitaxel and siRNA. Discover Applied Sciences, 7(8), 801.

60. Mohajeri S, et al. (2025). Targeted Gene Delivery to MCF-7 Cells via Polyspermine-PEG-Glucose/DNA Nanoparticles: Preparation and Characterization. Molecular Biotechnology, 1–19.

61. Mohajeri S, et al. (2025). Multifunctional magnetic nanocapsules for dual delivery of siRNA and chemotherapy to MCF-7 cells (Breast cancer cells). Naunyn-Schmiedeberg’s Archives of Pharmacology, 1–23.

62. Montoya-García C O, et al. (2023). Bioactive compounds of purslane (Portulaca oleracea L.) according to the production system: A review. Scientia horticulturae, 308, 111584.

63. Negrón L M, et al. (2025). Protonation-Induced Octamer–Hexadecamer Transition in Multiresponsive Supramolecular G-Quadruplexes.

64. Noruzpour M, et al. (2025). Delivery of Moringa oleifera extract via PLA-PEG-FA/chitosan-PLA NPs into breast cancer cell lines. BioNanoScience, 15(2), 287.

65. Noruzpuor M, et al. (2024). Green synthesis of metal nanoparticles using aqueous extract of Moringa oleifera L. and investigating their antioxidant and antibacterial properties. Applied Chemistry Today, 19(71), 283–302.

66. Ojah E O, et al. (2021). Phytochemical and antibacterial properties of root extracts from Portulaca oleracea Linn.(Purslane) utilised in the management of diseases in Nigeria. Journal of Medicinal Plants for Economic Development, 5(1), 103.

67. Oun R, et al. (2018). The side effects of platinum-based chemotherapy drugs: a review for chemists. Dalton transactions, 47(19), 6645–6653.

68. Purushothaman R, et al. (2024). Isolation and Identification of N-Hexadecanoic Acid from Excoecaria Agallocha L. and its Antibacterial and Antioxidant Activity. Journal of emerging technologies and innovative research, 11(1), 332–342.

69. Rahim M A, et al. (2023). Essential components from plant source oils: A review on extraction, detection, identification, and quantification. Molecules, 28(19), 6881.

70. Ramalho M J, et al. (2024). Folic-acid-conjugated poly (lactic-co-glycolic acid) nanoparticles loaded with gallic acid induce glioblastoma cell death by reactive-oxygen-species-induced stress. Polymers, 16(15), 2161.

71. Roopa M, et al. (2020). Comparative analysis of phytochemical constituents, free radical scavenging activity and GC-MS analysis of leaf and flower extract of Tithonia diversifolia (Hemsl.) A. Gray. International Journal of Pharmaceutical Sciences and Research, 11(10), 5081–5090.

72. Rubab M, et al. (2020). Bioactive Potential of 2-Methoxy-4-vinylphenol and Benzofuran from Brassica oleracea L. var. capitate f, rubra (Red Cabbage) on Oxidative and Microbiological Stability of Beef Meat. Foods, 9(5), 568.

73. Salim L, et al. (2020). Targeted delivery and enhanced gene-silencing activity of centrally modified folic acid–siRNA conjugates. Nucleic acids research, 48(1), 75–85.

74. Saygideger Y, et al. (2021). Antitumoral effects of Santolina chameacyparissus on non-small cell lung cancer cells. Journal of Experimental and Clinical Medicine, 38(3), 294–300.

75. Selvaraju R, et al. (2021). GC–MS and FTIR analysis of chemical compounds in Ocimum gratissimum plant. Biophysics, 66(3), 401–408.

76. Sh B, et al. (2024). Evaluation of antioxidant properties of essential oil, aqueous extract and metal nanoparticles biosynthesized from F. vulgare and their anticancer effect on two breast cancer cell lines (Sum-159, Hs-578T). Agricultural Biotechnology Journal, 16(1), 235–266.

77. Singh J, & Nayak, P. (2023). pH-responsive polymers for drug delivery: trends and opportunities. Journal of polymer science, 61(22), 2828–2850.

78. Sri P, et al. (2021). Antitumor Activity of Turnera subulata Sm.(Turneraceae) in Hep G2 Cancer Cell Line. Journal of Pharmaceutical Research International, 33(29A), 191–199.

79. Srivastava R, et al. (2023). Multipurpose benefits of an underexplored species purslane (Portulaca oleracea L.): A critical review. Environmental Management, 72(2), 309–320.

80. Subroto E, et al. (2023). Solid Lipid Nanoparticles: Review of the Current Research on Encapsulation and Delivery Systems for Active and Antioxidant Compounds. Antioxidants, 12(3), 633.

81. Swetha T A, et al. (2023). A comprehensive review on polylactic acid (PLA)–Synthesis, processing and application in food packaging. International Journal of Biological Macromolecules, 123715.

82. Tadigiri S, et al. (2020). Isolation and characterization of chemical constituents from B. amyloliquefaciens and their nematicidal activity. Mortality, 8(12h), 24h.

83. Thuy B T P, et al. (2021). Screening for Streptococcus pyogenes antibacterial and Candida albicans antifungal bioactivities of organic compounds in natural essential oils of Piper betle L., Cleistocalyx operculatus L. and Ageratum conyzoides L. Chemical Papers, 75, 1507–1519.

84. Tian R, et al. (2022). Analysis of aromatic components of two edible mushrooms, Phlebopus portentosus and Cantharellus yunnanensis using HS-SPME/GC-MS. Results in Chemistry, 4, 100282.

85. Tleubayeva M I, et al. (2021). Component composition and antimicrobial activity of CO2 extract of Portulaca oleracea, growing in the territory of Kazakhstan. The Scientific World Journal, 2021(1), 5434525.

86. Tomou E-M, et al. (2023). Recent advances in nanoformulations for quercetin delivery. Pharmaceutics, 15(6), 1656.

87. Tsimogiannis D, et al. (2017). DPPH radical scavenging and mixture effects of plant o-diphenols and essential oil constituents. European journal of lipid science and technology, 119(9), 16003473.

88. Tuan N H, et al. (2021). Chemical composition and antibacterial properties of essential oil extracted from the leaves and the rhizomes of Stahlianthus thorelii Gagnep.(Zingiberaceae). Journal of Essential Oil Bearing Plants, 24(6), 1365–1372.

89. Tune B X J, et al. (2022). Matrix metalloproteinases in chemoresistance: regulatory roles, molecular interactions, and potential inhibitors. Journal of oncology, 2022(1), 3249766.

90. Verkhovskii R A, et al. (2023). Current principles, challenges, and new metrics in pH-responsive drug delivery systems for systemic cancer therapy. Pharmaceutics, 15(5), 1566.

91. Visan A I, et al. (2021). Degradation behavior of polymers used as coating materials for drug delivery—A basic review. Polymers, 13(8), 1272.

92. Wadhawan A, et al. (2022). Anticancer biosurfactant-loaded PLA–PEG nanoparticles induce apoptosis in human MDA-MB-231 breast cancer cells. ACS omega, 7(6), 5231–5241.

93. Wang Y, et al. (2021). Risk assessment of agricultural plastic films based on release kinetics of phthalate acid esters. Environmental science & technology, 55(6), 3676–3685.

94. Xia W, et al. (2021). Targeted delivery of drugs and genes using polymer nanocarriers for cancer therapy. International journal of molecular sciences, 22(17), 9118.

95. Xie Y, et al. (2022). A new strategy based on PCA for inter-batches quality consistency evaluation. Journal of Pharmaceutical and Biomedical Analysis, 217, 114838.

96. Yang T-Y, et al. (2012). Delayed foreign body reaction after fixation of distal radius fracture with biodegradable implant. Formosan Journal of Musculoskeletal Disorders, 3(1), 27–30.

97. Yun H J, et al. (2022). Induction of cell cycle arrest, apoptosis, and reducing the expression of MCM proteins in human lung carcinoma A549 cells by cedrol, isolated from Juniperus chinensis. Journal of Microbiology and Biotechnology, 32(7), 918.

98. Zhang D, et al. (2024). Effect of volatile compounds produced by Weissella cibaria BWL4 on Botrytis cinerea infection in fruit and complete genome sequence analysis of BWL4. Postharvest Biology and Technology, 213, 112917.

99. Zhang Q-Y, et al. (2018). Natural product interventions for chemotherapy and radiotherapy-induced side effects. Frontiers in pharmacology, 9, 1253.

100. Zhang Y, et al. (2025). Individual and combined effects of ZnO nanoparticles and cinnamon essential oil on Salmonella typhimurium in milk-based beverage. European Food Research and Technology, 1–10.

101. Zhao T, et al. (2024). Chemical characterization, antioxidant, antimicrobial, enzyme inhibitory and cytotoxic activities of Illicium lanceolatum essential oils. Arabian Journal of Chemistry, 17(1), 105366.

102. Zhao Y, et al. (2023). Cedrol, a Major Component of Cedarwood Oil, Ameliorates High-Fat Diet-Induced Obesity in Mice. Molecular Nutrition & Food Research, 67(14), 2200665.

103. Zhou F, et al. (2023). The commonly used stabilizers for phytochemical-based nanoparticles: stabilization effects, mechanisms, and applications. Nutrients, 15(18), 3881.

104. Zhu H, et al. (2010). Identification of Portulaca oleracea L. from different sources using GC– MS and FT-IR spectroscopy. Talanta, 81(1-2), 129–135.

